# Identification of a conserved neutralizing epitope present on spike proteins from all highly pathogenic coronaviruses

**DOI:** 10.1101/2021.01.31.428824

**Authors:** Yimin Huang, Annalee W. Nguyen, Ching-Lin Hsieh, Rui Silva, Oladimeji S. Olaluwoye, Rebecca E. Wilen, Tamer S. Kaoud, Laura R. Azouz, Ahlam N. Qerqez, Kevin C. Le, Amanda L. Bohanon, Andrea M. DiVenere, Yutong Liu, Alison G. Lee, Dzifa Amengor, Sophie R. Shoemaker, Shawn M. Costello, Susan Marqusee, Kevin N. Dalby, Sheena D’Arcy, Jason S. McLellan, Jennifer A. Maynard

**Affiliations:** Department of Molecular Biosciences, The University of Texas, Austin, TX 78712, USA; Department of Chemical Engineering, The University of Texas, Austin, TX 78712, USA; Division of Chemical Biology and Medicinal Chemistry, The University of Texas, Austin, TX 78712, USA; Department of Chemistry and Biochemistry, The University of Texas, Dallas, TX 75080, USA; Department of Molecular & Cell Biology, University of California, Berkeley, CA 94720, USA; Biophysics Graduate Program, University of California, Berkeley, CA 94720, USA; Department of Chemistry, University of California, Berkeley, CA 94720, USA; LaMontagne Center for Infectious Diseases, The University of Texas, Austin, TX 78712, USA

## Abstract

Three pathogenic human coronaviruses have emerged within the last 20 years, with SARS-CoV-2 causing a global pandemic. Although therapeutic antibodies targeting the SARS-CoV-2 spike currently focus on the poorly conserved receptor-binding domain, targeting essential neutralizing epitopes on the more conserved S2 domain may provide broader protection. We report an antibody binding an epitope conserved in the pre-fusion core of MERS-CoV, SARS-CoV and SARS-CoV-2 spike S2 domains. Antibody 3A3 binds a conformational epitope with ~2.5 nM affinity and neutralizes spike from SARS-CoV, SARS-CoV-2 and variants of concern in *in vitro* pseudovirus assays. Hydrogen-deuterium exchange mass spectrometry identified residues 980-1006 in the flexible hinge region at the S2 apex as the 3A3 epitope, suggesting 3A3 prevents the S2 conformational rearrangements required for conversion to the spike post-fusion state and virus-host cell fusion. This work defines a conserved vulnerable site on the SARS-CoV-2 S2 domain and guides the design of pan-protective spike immunogens.

## INTRODUCTION

COVID-19, the disease caused by severe acute respiratory syndrome coronavirus 2 (SARS-CoV-2), has been responsible for over 4 million deaths worldwide since it was identified in late 2019. This pandemic is the latest and largest of three deadly coronavirus outbreaks, including those caused by SARS-CoV in 2002 and Middle East respiratory syndrome coronavirus (MERS-CoV) in 2012. Although COVID-19 has a case fatality rate estimated at ~1– 3% versus ~10% for SARS-CoV and ~34% for MERS^1^, it has proven to be far more infectious and its spread has impacted every aspect of society worldwide. Four other coronaviruses are known to infect humans, resulting in relatively mild upper respiratory disease and symptoms: 229E, OC43 (both discovered in the 1960s), NL63 (2004), and HKU1 (2005). All seven coronaviruses are zoonotic, and many other coronaviruses are endemic in animals, foreshadowing future coronavirus outbreaks.

Coronaviruses are enveloped positive sense, single-stranded RNA viruses that invade target cells by fusion of the viral envelope with the target cell membrane, mediated by the spike glycoprotein. The spike is a homo-trimer comprised of S1 and S2 domains: S2 forms a stalk-like structure proximal to the viral envelope while S1 forms a cap covering the S2 apex. Each S1 domain monomer contains an N-terminal domain (NTD) and a receptor-binding domain (RBD) that overlaps adjacent NTDs and RBDs within the trimer, forming a responsive surface that allows each RBD to extend to the “up” position for receptor binding or tuck into the “down” position for immune shielding. When RBDs engage a receptor in the up position and target-cell-anchored proteases prime the spike, the S1 domain is released from S2, propelling the fusion peptide into the target cell surface. Simultaneously, S2 undergoes a massive structural rearrangement to bring the viral envelope into contact with the target cell, initiating fusion and leaving spike in the post-fusion state^2^.

A powerful strategy to prevent coronavirus fusion is to disrupt the interaction between the RBD and its host cell receptor. For SARS-CoV-2, the receptor is angiotensin-converting enzyme 2 (ACE2), and antibodies that block RBD binding to ACE2 are both potently neutralizing and common in convalescent patient sera^3^. Accordingly, most monoclonal antibody therapies in development target this interaction^4^. However, RBD sequences from different coronaviruses vary considerably to allow for binding to different host cell receptors. Even within the three human-relevant coronaviruses that bind ACE2 (SARS-CoV, SARS-CoV-2 and NL63), RBD-binding antibodies exhibit limited cross-reactivity^5^, consistent with the low level of S1 sequence conservation (~20–24% identity, ~41–52% similarity; Supplementary Fig. 1). As a result, it has been difficult to identify antibodies binding multiple lineage B β-coronaviruses, such as SARS-CoV and SARS-CoV-2^6^, much less to include the lineage C MERS-CoV. In the CoV-AbDab database^7^ there are only currently two antibodies reported to neutralize SARS-CoV, SARS-CoV-2 and MERS-CoV and neither bind the S1 domain.

By contrast, S2 is the most conserved spike domain, with 63–98% sequence similarity in pairwise comparisons across the seven human coronaviruses (Supplementary Fig. 1). Moreover, the functionally analogous domain in the fusion proteins from influenza virus, respiratory syncytial virus (RSV) and human immunodeficiency virus (HIV) contains potently neutralizing epitopes^8,9^, prompting speculation that the spike S2 domain may also serve as an effective target for neutralizing antibodies. Multiple mechanisms of spike fusion inhibition are being explored, including those that target the S2 domain to prevent S2 conformational rearrangement^10,11^, block the fusion peptide^12–14^, and interfere with S2 proteolytic processing^15^. Strategies targeting the S2 domain aim to induce premature S1 shedding and non-productive transformation into the post-fusion spike. Antibodies binding the S2 domain of SARS-CoV-2 or conserved epitopes on S2 are critical to support these efforts, but few have been reported in detail. Structures of S2-binding antibodies are limited to two peptide-antibody complexes for antibodies that bind the membrane-proximal stem helix of the spike^10,11^. Of the over 2500 anti-SARS-CoV-2 spike antibodies reported in the CoV-AbDab database, less than 3% are reported to bind both SARS-CoV and SARS-CoV-2 spike and less than 10% bind the S2 region. A total of 13 reported S2 binding antibodies have neutralizing capacity^7^.

Here, we aimed to identify unique epitopes conserved across all known highly pathogenic coronaviruses. To focus immune responses on the S2 domain, we immunized mice with a stabilized MERS-CoV S2 protein and isolated antibodies using alternate rounds of selection with the MERS-CoV S2 and intact SARS-CoV-2 spike using phage display. Of three high affinity, cross-reactive antibodies characterized, 3A3 neutralizes SARS-CoV-2 spike in cell fusion and pseudovirus assays by binding a conformational epitope at the apex of the S2 domain. This epitope is available for 3A3 binding only when the spike protein is fully open or splayed into three C-terminally tethered protomers in a recently described spike conformation^16^. The 3A3 epitope defines a novel conserved site of vulnerability, which is increasingly relevant for more transmissible spike variants, future pan-coronavirus vaccine designs and other therapeutic strategies.

## RESULTS

### Isolation of antibodies binding the S2 domain of SARS-CoV, SARS-CoV-2 and MERS-CoV spikes

Three unique antibody families binding a broad range of spike variants were identified from a mouse immune phage library. Balb/c mice were immunized with stabilized MERS-CoV S2 protein and boosted four weeks later, resulting in robust serum antibody titers (detectable at >1:10,000 dilution). The MERS-CoV S2 protein includes amino acid residues 763–1291 of the MERS-CoV spike protein with a C-terminal T4 phage fibritin (foldon) domain that assembles into a pre-fusion trimer. In this work, “spike” refers to the extracellular coronavirus fusogen domains containing homologous 2P mutations (residues 986 and 987^17^ in SARS-CoV-2) C-terminally fused to a foldon domain, whereas “unstabilized” refers to spike variants as expressed on the virion without stabilizing changes, with other variations noted. “Wild-type” SARS-CoV-2 spike refers to the spike sequence originally reported in January of 2020 for 19A (Wuhan-Hu-1) SARS-CoV-2.

To generate immune antibody libraries, total mRNA was isolated from the mouse spleens and reverse transcribed; the antibody variable regions were then amplified^18^ and inserted into the pMopac24 M13 bacteriophage display vector^19^ to express scFv-c-myc tag-pIII fusion proteins. The phage display library (~3.1×10^8^ individual clones) was subjected to four rounds of panning with immobilized antigen: anti-c-myc antibody to deplete truncated non-expressing clones, then MERS-CoV S2, and finally SARS-CoV-2 spike at high followed by moderate coating concentrations. Phage clones isolated after rounds 3 and 4 were confirmed to bind MERS-CoV S2 and SARS-CoV-2 spike by ELISA and evaluated for binding to uncoated plates and the foldon domain. While no clones exhibited plate or milk binding, ~85% of round 3 and 4 clones bound the shared foldon domain fused to the RSV F protein^20^ (Supplementary Fig. 2). One of these foldon binders, 3E11, was carried forward as a control antibody, while the remaining clones identified three spike-specific families: 3A3, 4A5, and 4H2. Antibody 3A3 and close relatives appeared in round 3 and were enriched to ~10% of the population after round 4, while 4A5 and 4H2 were unique sequences isolated after round 4 of panning.

### Cross-reactive antibodies bind the spike S2 domain with high affinity and specificity

After expression in ExpiCHO cells and purification of 3A3, 4A5, 4H2, and 3E11 as full-length antibodies with human IgG1 and kappa constant domains, the antibodies were biophysically characterized to measure spike binding affinity by biolayer interferometry (BLI) and surface plasmon resonance (SPR), thermal unfolding by thermal shift, polydispersity by size exclusion chromatography (SEC), and purity by SDS-PAGE (Table 1 and Supplementary Figs. 3 and 4). The antibodies all appeared as expected for intact immunoglobulins, with 4A5 exhibiting a slightly delayed SEC retention volume. All four antibodies exhibited low- to mid- nanomolar equilibrium affinities for the SARS-CoV-2 spike^17^ as measured by BLI. Additional measurements with SARS-CoV-2 HexaPro, an ultra-stable SARS-CoV-2 spike variant with six proline substitutions relative to wild-type spike^21^, confirmed these affinities. The binding of 3A3 Fab to the isolated S2 domain of SARS-CoV-2 HexaPro revealed similar ~2.5 nM K_d_ values by both SPR and BLI. The on- and off-rate constants for HexaPro S2 binding in the absence of S1 were ~2×10^6^ M^−1^s^−1^ and 5×10^−3^ s^−1^, respectively (Supplementary Fig. 4).

**Table 1.** Antibody binding affinities for spike variants†.

|  | Spike variant K<sub>d</sub> (nM) |  |  |  |  |  |  |
| --- | --- | --- | --- | --- | --- | --- | --- |
|  | SARS-2 | HexaPro<br>S2 | HexaPro | HexaPro<br>aglyc | SARS-1 | MERS | MERS<br>S2 |
| **3A3** | 2.7 ± 0.3 | 3.2 ± 0.3 | 8 ± 3 | 7.9 ± 0.8 | 20 ± 2 | 23 ± 1.2 | 5.9 ± 0.6 |
| **4A5** | 17 ± 4 | 11.0 ± 0.1 | 4 ± 1 | 10 ± 2 | 4 ± 0.9 | ND | ND |
| **4H2** | 29 ± 4 | 63 ± 7 | 5.8 ± 0.9 | 9 ± 2 | 20 ± 2 | ND | ND |
| **3E11** | 5.5 ± 0.7 | 8 ± 4 | 3.5 ± 0.6 | 11 ± 2 | 5.7 ± 0.5 | ND | ND |

† K<sub>d</sub> values represent the equilibrium affinity for full-length IgGs immobilized on anti-Fc tips capturing the indicated SARS-2 spike or spike domain, as measured by BLI with standard error shown. All fits had R<sup>2</sup>>0.90.
ND = no data.

To assess the phylogenetic range of spike recognition by these three antibodies, binding to SARS-CoV, SARS-CoV-2, MERS-CoV, and HKU1 as well as SARS-CoV-2 HexaPro spikes, and RSV F-foldon control was assessed by ELISA (Fig. 1a and Supplementary Fig. 5) and validated with BLI K_d_ measurements (Table 1 and Supplementary Fig. 4d). Antibodies 3A3, 4A5 and 4H2 retained strong binding to the SARS-CoV spike, MERS-CoV spike and MERS-CoV S2 domain but exhibited significantly reduced binding to HKU1 spike and no detectable binding to RSV F-foldon. Specifically, 3A3 exhibited an affinity of 20 nM for SARS-CoV and 23 nM for MERS-CoV intact spikes. The foldon binding antibody 3E11 showed relatively high binding affinity ~3–11 nM across all spikes and the RSV F-foldon as expected since these proteins all include foldon domains for stabilization.

**Fig. 1.**
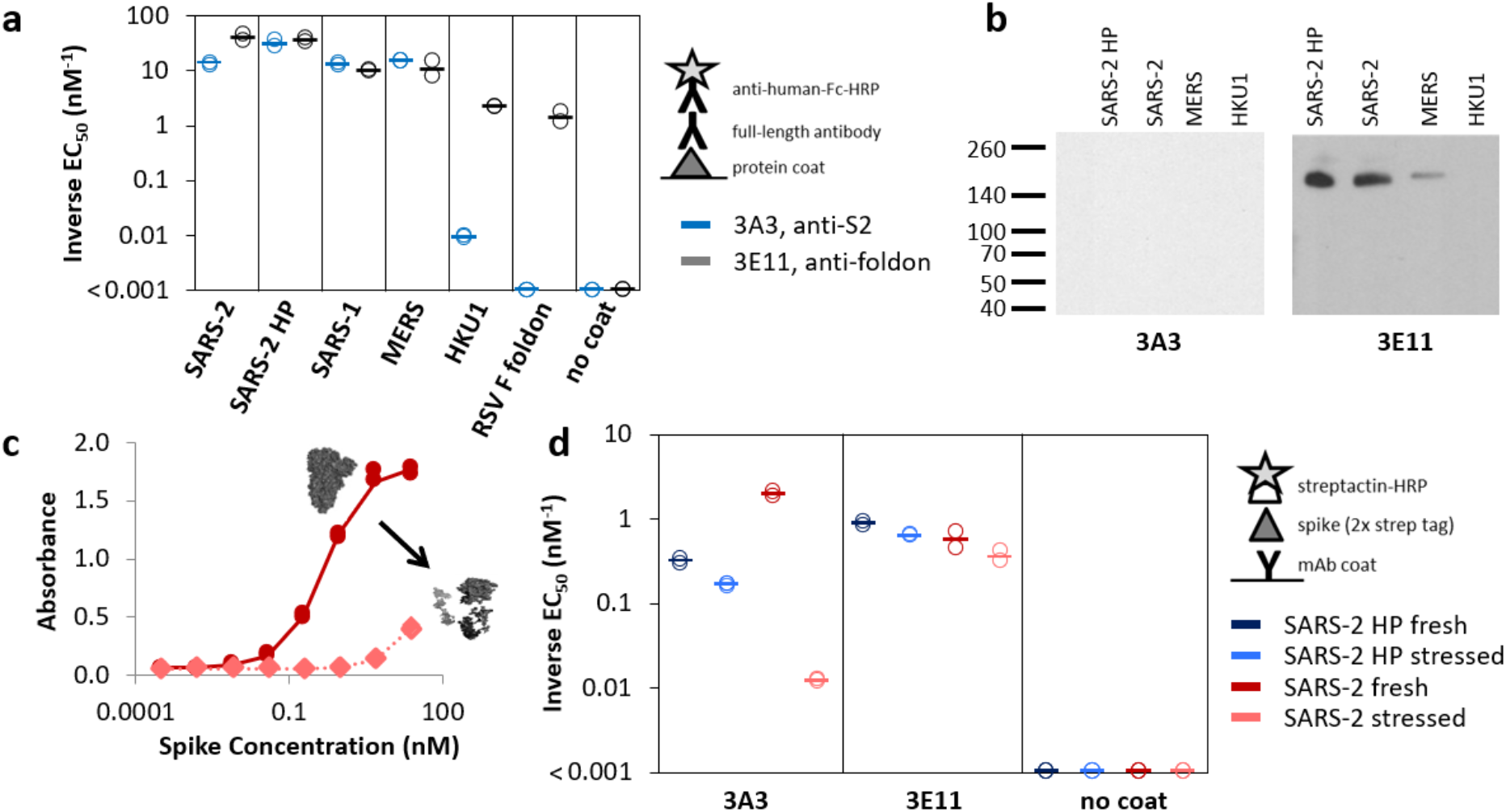
Antibody 3A3 binds to SARS-CoV and MERS-CoV spike proteins. (**a**) Full-length anti-S2 antibody 3A3 (blue) and anti-foldon antibody 3E11 (grey) were tested for binding to SARS-CoV-2 (SARS-2), SARS-CoV-2 HexaPro (SARS-2 HP), SARS-CoV (SARS-1), MERS-CoV, HKU1, RSV F foldon coated, or milk (no coat) proteins by ELISA. **b** 3E11 binds reduced, denatured SARS-2 HP, SARS-2, and MERS-CoV spike proteins by western blot, but 3A3 does not. Neither antibody bound to HKU1 spike by western blot. The ladder molecular weight is labeled in kDa on the left side. **c** ELISA capture of fresh (red circles) or stressed (pink diamonds) SARS-2 spike on 3A3 coated plates. **d** Antibodies coated on ELISA plates captured fresh (dark blue or red) or stressed (pink or light blue) SARS-2 HP (blue) or SARS-2 (red) spike proteins. For both **a** and **d**, duplicate dilutions of spike over ~5 log in concentration were used to calculate EC_50_ values. For dilution series in which no binding was observed, EC_50_ was assumed to be >1000 nM. Open symbols are replicate data and filled rectangles are average data.

### Antibody 3A3 binds a conformationally sensitive S2 epitope

To further investigate the basis for spike protein recognition, the SARS-CoV-2, SARS-CoV-2 HexaPro, MERS-CoV, and HKU1 spikes were fully denatured and reduced and subjected to western blotting. Quadruplicate blots were incubated with each full-length antibody and binding to the spike polypeptide was detected with anti-human-Fc-HRP and chemiluminescent peroxidase substrate (Fig. 1b and Supplementary Fig. 5c). Antibodies 4A5, 4H2, and 3E11 detected linearized SARS-CoV-2, SARS-CoV-2 HexaPro, and MERS-CoV spike proteins, with undetectable binding to HKU1 spike. In contrast, 3A3 did not bind any spike protein in western blot, indicating that the 3A3 antibody binds a conformational epitope, while the other three antibodies recognize epitopes with significant linear components.

To better understand the 3A3 conformational epitope, we evaluated 3A3 binding to fresh versus stressed SARS-CoV-2 spike by ELISA (Fig. 1c and d, and Supplementary Fig. 5d). Soluble SARS-CoV-2 spike is sensitive to freeze-thaw stresses; by contrast, SARS-CoV-2 HexaPro includes four additional proline substitutions and exhibits resistance to freeze-thaw stresses^21^. We stressed spike by three freeze/thaw cycles, which produced SDS-PAGE-detectable aggregates in the SARS-CoV-2 spike but not SARS-CoV-2 HexaPro (Supplementary Fig. 6) and captured spike on antibody-coated ELISA plates. Antibodies 4A5, 4H2, and 3E11 bound fresh and stressed SARS-CoV-2 spike proteins similarly, with 4A5 and 4H2 binding stressed spike slightly better (Supplementary Fig. 5d). However, 3A3 bound stressed 2P spike with a ~150-fold worse EC_50_, while binding of stressed SARS-CoV-2 HexaPro was unaffected, indicating a reduced capacity for 3A3 to bind misfolded and/or aggregated spike.

### Antibody 3A3 neutralizes spike in *in vitro* cellular fusion and pseudovirus infection assays

To investigate the ability of 3A3, 4A5, and 4H2 to impact spike function, we first employed a mammalian cell fusion assay (Fig. 2a and b, and Supplementary Fig. 7). A CHO cell line expressing wild-type SARS-CoV-2 spike and GFP was incubated with ACE2-expressing HEK 293 cells dyed with the red fluorescent Cell Trace Far Red stain. After 24 hours, large syncytia formed with green CHO cell fluorescence overlapping ~70% of red HEK 293 cell fluorescence, indicating fusion of the CHO and HEK 293 membranes in the presence of no antibody (Supplementary Fig. 7a) or 670 nM irrelevant human IgG1 (Supplementary Fig. 7d), 4A5 or 4H2 antibodies (Supplementary Fig. 8). Minimal fluorescence colocalization occurred if either the CHO cells did not express SARS-CoV-2 spike (Supplementary Fig. 7b) or the HEK 293 cells did not express ACE2 (Supplementary Fig. 7c). Incubation with 670 nM or 67 nM of 3A3 significantly reduced colocalization to ~50% (p <0.0001; Fig. 2a). While 6.7 nM 3A3 had minimal impact on fluorescence colocalization (Supplementary Fig. 7e), analysis of average cell size showed significantly reduced syncytia size in the presence of as little as 6.7 nM 3A3 (Fig. 2b).

**Figure 2.**
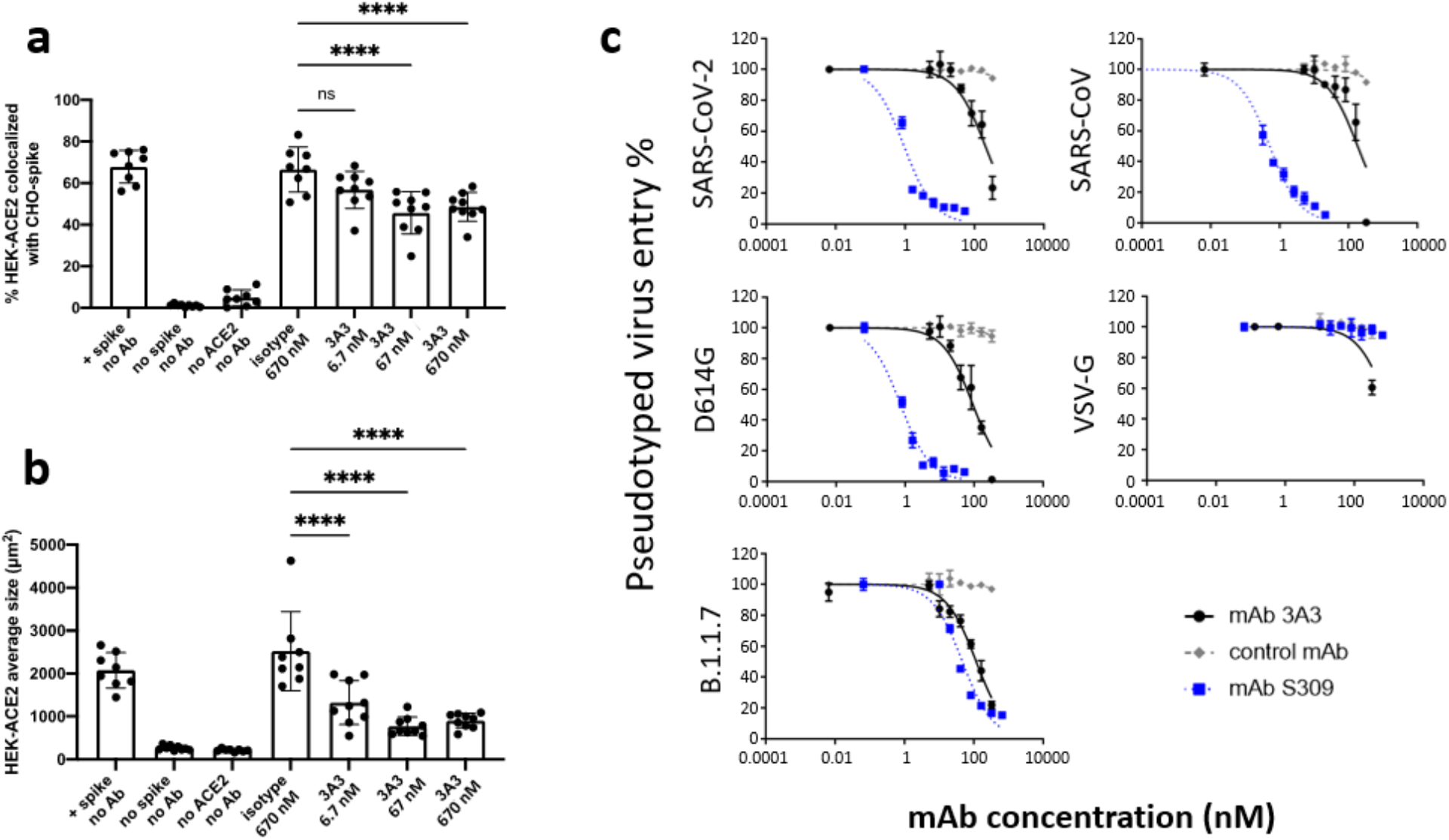
Antibody 3A3 binds and inhibits the fusion activity of unstabilized spike in both a mammalian fusion assay and pseudovirus neutralization assay. **a,b** HEK 293 cells stably expressing surface ACE2 (HEK-ACE2) were stained with Cell Trace Far Red and incubated with a CHO-based cell line transiently expressing unstabilized, wild-type SARS-CoV-2 spike and EGFP and preincubated with 3A3 antibody or isotype control. **a** The percentage of HEK-ACE2 pixels (red) colocalizing with spike expressing CHO pixels (green) was analyzed with the JACoP plugin for ImageJ. **b** The same images were analyzed for the average cell size of fused HEK-ACE2 with ImageJ as a second statistical method to test the cell fusion level. Shown are the mean and standard deviation of at least 160 cells per condition from 8– 9 independent images. The statistical analysis of average cell sizes under different conditions were performed with ANOVA followed by Tukey’s HSD test. Results shown are representative of four independent experiments; **** p<0.0001. **c** Neutralization assays were performed with HIV particles pseudotyped with unstabilized SARS-CoV-2 wild-type spike (SARS-2 spike, filled circles), the SARS-CoV-2 D614G mutant (D614G spike, triangles), B.1.1.7 SARS-CoV-2 (variant alpha) spike (B.1.1.7 spike, diamonds) and SARS-CoV spike (SARS-1 spike, squares). The pseudoviruses were incubated with different doses of each antibody (3A3, black; S309, blue; or an isotype control with irrelevant binding, grey) for 1 hour at room temperature before adding to HEK 293T cells stably expressing ACE2. Viral entry was detected by luciferase luminescence 60–72 hrs later. The entry efficiency of SARS-CoV-2 pseudoviruses without any treatment was considered 100%. Treatment of every pseudovirus with S309 or 3A3 resulted in significant (p<0.01) neutralization versus isotype control antibody with matched virus. Statistical analyses of neutralization IC_50_ were performed with ANOVA followed by Tukey’s HSD test.

We next assessed 3A3 neutralization in an *in vitro* pseudovirus neutralization assay. Antibody 3A3, potently neutralizing antibody S309^22^, or an isotype control antibody was preincubated for one hour with pseudotyped lentivirus expressing SARS-CoV spike, wild-type SARS-CoV-2 spike, SARS-CoV-2 D614G spike, or SARS-CoV-2 B.1.1.7 (variant Alpha) spike, all without stabilizing modifications, then added to HEK 293 cells expressing ACE2. The pseudovirus induced luciferase expression in infected cells and the extent of infection was tracked for 72 hours (Fig. 2c). With one hour of preincubation, the positive control S309 had an IC_50_ of ~1 nM against SARS-CoV and SARS-CoV-2 spike, similar to values found with MLV^22^ pseudovirus. Antibody S309 was less potent against the B.1.1.7 pseudotyped-virus than wild-type or D614G, as previously reported^23^, while the isotype control had no effect on any pseudoviruses. By contrast, 3A3 blocked infection of wild-type spike pseudovirus with an IC_50_ of 212 nM, D614G spike pseudovirus with an IC_50_ of 94 nM, and B.1.1.7 spike pseudovirus with an IC_50_ of 119 nM (Table 2). SARS-CoV spike pseudotyped virus was neutralized by 3A3 with an IC_50_ of 188 nM. Consistent with pseudovirus neutralization, 3A3 bound unstabilized D614G spike displayed on mammalian cells more readily than wild-type spike by flow cytometry (Supplementary Fig. 9).

**Table 2.** Antibody neutralization of pseudovirus IC_50_ values.

|  | Pseudovirus IC<sub>50</sub><br>(95% confidence interval), nM |  |  |  |  |
| --- | --- | --- | --- | --- | --- |
|  | SARS-2 | D614G | B.1.1.7 | SARS-1 | VSV-G |
| **3A3** | 212<br>(167-273) | 94<br>(73-122) | 119<br>(103-138) | 188<br>(127-283) | 924<br>(668-1380) |
| **S309** | 1.0<br>(0.8-1.2) | 0.8<br>(0.6-0.9) | 48<br>(39-59) | 0.5<br>(0.5-0.6) | >5000 |
| **Isotype**<br>**control** | >5000 | >5000 | >5000 | >5000 | >5000 |

### The neutralizing 3A3 epitope is located on the S2 hinge

To identify the specific epitope recognized by 3A3, we used hydrogen-deuterium exchange mass spectrometry (HDX). We measured deuterium uptake of the SARS-CoV-2 HexaPro spike alone, as well as bound by the 3A3 IgG or 3A3 Fab (Supplementary Tables 1 and 2). Complexes were formed with excess antibodies such that the SARS-CoV-2 HexaPro spike was ~90% bound in both cases. We tracked 192 unmodified peptides through the deuteration time course (10^1^, 10^2^, 10^3^, and 10^4^ s), and did not search for glycosylated peptides as de-glycosylation had a small effect on 3A3 affinity (Table 1 and Supplementary Fig. 10). Analysis of the raw deuterium uptake in the SARS-CoV-2 HexaPro spike alone is consistent with a trimer during exchange; with relatively low deuterium uptake in the helix at the center of the trimer and high deuterium uptake in the HR1 helix at the trimer’s surface (Supplementary Fig. 11). Further analysis of the isotopic distribution width of peptides from regions of spike reported to display bimodality^16^ is consistent with the trimeric spike sampling an open conformation (Supplementary Table 3).

Antibody epitopes were identified by examining the difference in deuterium uptake between SARS-CoV-2 HexaPro spike in the free and antibody-bound states. We defined a significant difference in a change in deuterium uptake greater than 0.2 Da with a p-value less than 0.01 (Fig. 3a and b). The binding of 3A3 IgG caused a significant decrease in 12 peptides that redundantly span residues 980 to 1006 of the SARS-CoV-2 HexaPro spike (Fig. 3c and Supplementary Fig. 12a). These peptides have reduced deuterium uptake with 3A3 IgG at several timepoints during the exchange reaction. An identical result was obtained with the 3A3 Fab (Fig. 3b and c and Supplementary Fig. 12). In contrast to 3A3 IgG and Fab, similar experiments with 4A5 and 4H2 antibodies showed no significant difference in deuterium uptake upon antibody addition (Supplementary Fig. 13). The epitopes recognized by 4A5 and 4H2 are distinct from 3A3 and possibly lie in regions where we lack peptides. Taken altogether, these data show that the 3A3 epitope lies within residues 980 to 1006 of the SARS-CoV-2 HexaPro spike.

**Fig. 3.**
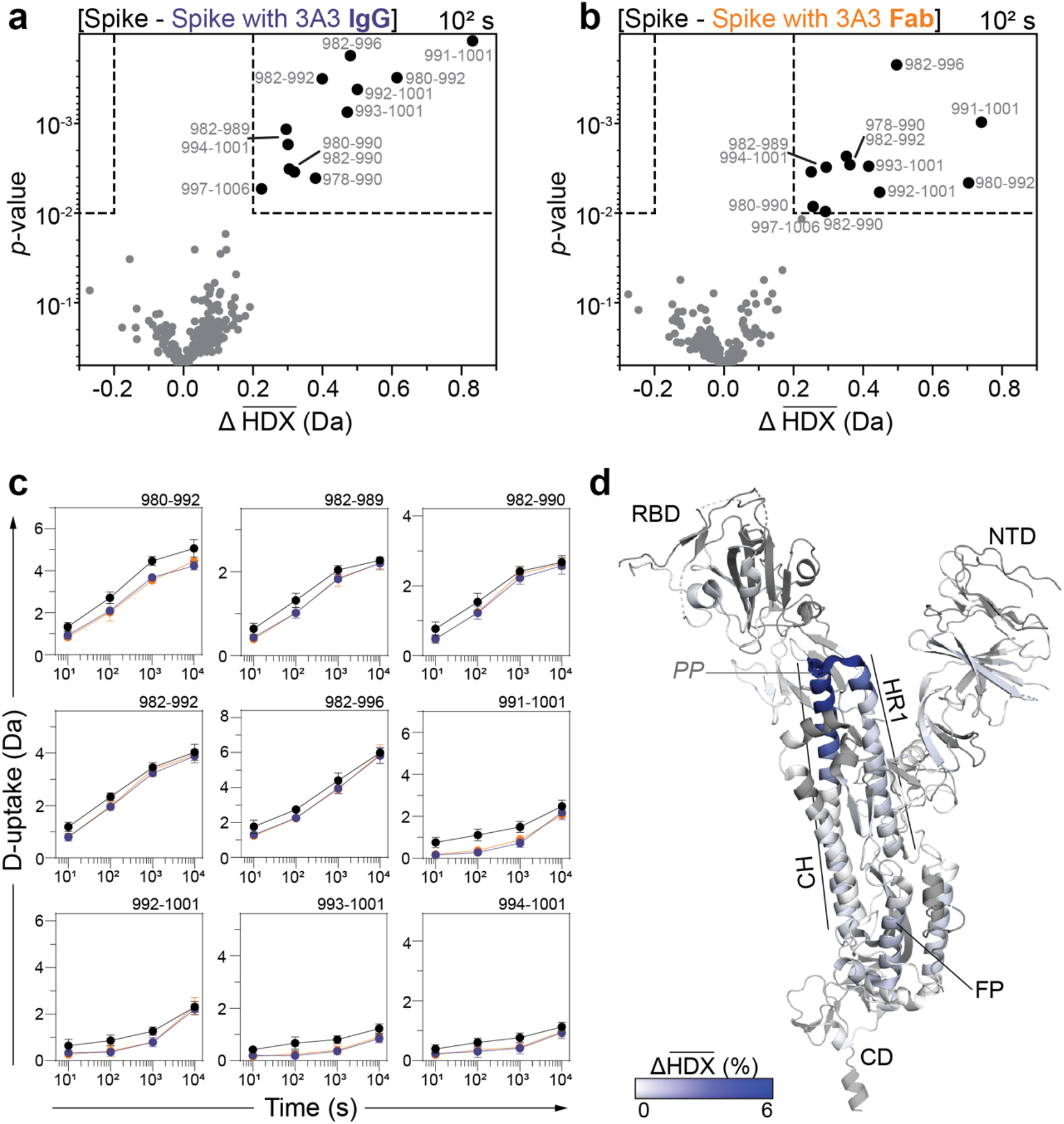
The 3A3 epitope is located at the apex of the S2 domain. Volcano plots showing changes in deuterium uptake in SARS-2 HexaPro spike peptides upon addition of 3A3 IgG **a** or Fab **b** after 10^2^ s exchange. Significance cutoffs are an average change in deuterium uptake greater than 0.2 Da and a *p*-value less than 10^−2^ in a Welch’s t-test (hatched box). Black dots indicate peptides with a significant decrease in deuterium uptake and their boundaries are labeled. **c** Deuterium uptake plots for peptides with a significant decrease in deuterium uptake upon addition of 3A3 (see Supplementary Fig. 12). Traces are SARS-2 HexaPro spike alone (black), with 3A3 IgG (blue), and with 3A3 Fab (orange). Error bars are ±2σ from 3 or 4 technical replicates. Y-axis is 70% of max deuterium uptake assuming the N-terminal residue undergoes complete back-exchange. Data have not been corrected for back-exchange. **d** Monomeric SARS-CoV-2 2P spike (PDB: 6VSB chain B) colored according to the difference in deuterium fractional uptake between SARS-2 HexaPro spike alone and with 3A3 IgG at 10^2^ s exchange. The figure was prepared using DynamX per residue output without statistics and Pymol. Residues lacking coverage are indicated in grey. Structural features are labeled, including the 2P mutations at residues 986 and 987 (shown as sticks).

Mapping the difference in deuterium uptake between free and 3A3-bound states onto the structure localizes the epitope to the apex of the S2 domain, distal to the viral envelope (Fig. 3d). It covers the end of the HR1 helix, the two stabilizing proline mutations (residues 986 and 987) and the beginning of the CH helix. This region is highly conserved in sequence and structure across all β-coronaviruses known to infect humans (Fig. 4, a and b), with Cα atoms RMSD ranging from 0.6 Å for HKU1 to 3.1 Å for MERS-CoV. Fourteen of the solvent exposed residues within the epitope were altered to assess the impact on 3A3 binding. Three, S982A (present in the B.1.1.7 SARS-CoV-2 Alpha variant), L984V and R995A improved 3A3 binding by ELISA, while five (D985L, E988Q or I, D994A, L1001A and Q1002A) significantly reduced 3A3 binding (Fig. 4c and d, Supplementary Fig. 15). The most substantial impact was the near ablation of binding by the substitutions at positions D985 and E988. These residues form a negatively charged patch adjacent to the 2P changes, P986 and P987. Since E988Q is present in the spike proteins of the α-coronaviruses NL63 and 229E, these data suggest 3A3 neutralization may be limited to β-coronaviruses. In addition to the clear role of residues near the S2 hinge, residues D994, L1001 and Q1002 lie deeper in the S2 core and are hidden in prefusion trimer structures^17,21^, yet appear to contribute to 3A3 spike recognition. Substitutions that improve binding are at the protomer interfaces and may destabilize the closed trimer conformation. Confirming this epitope in MERS-CoV spike, 3A3 strongly bound MERS-CoV S2 but not a mutant with the apex of the S2 domain replaced with a linker (Supplementary Figure 13).

**Figure 4.**
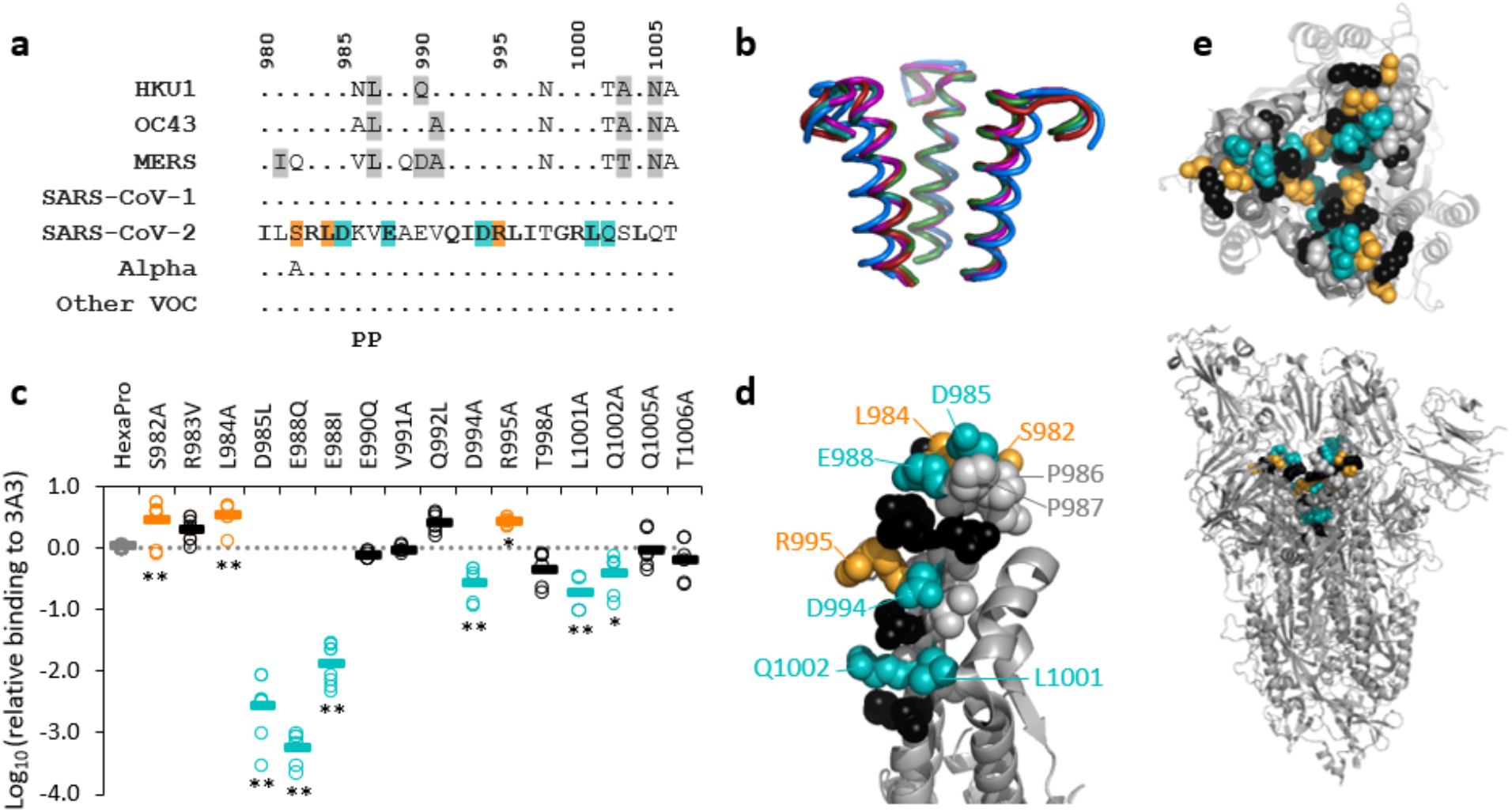
Structural location of 3A3 epitope and implications for antibody binding. The 3A3 epitope (SARS-CoV-2 amino acids 980–1006) is highly conserved across the spike a sequences and b structures of beta coronaviruses known to infect humans, including B.1.1.7 (Alpha) and the Beta, Gamma, Delta, and Epsilon variants of concern (Other VOC). In **a**, identical residues are indicated by a dot and similar residues are highlighted in grey. Residues conserved across all listed β-coronaviruses are in bold in the SARS-CoV-2 epitope. Residues that lost binding to 3A3 when altered as shown in **c** are in teal highlight and those whose disruption improved binding are orange. The location of the two proline mutations introduced to 2P variants are shown below the alignment. In **b**, the structure of each epitope is displayed as follows: SARS-2 (6VSB) – red, SARS-1 (6CRV, RMSD = 0.8 Å) – magenta, MERS-CoV (5X5C, RMSD = 3.1 Å) – blue, HKU1 (5I08, RMSD = 0.5 Å) – teal, OC43 (6OHW, RMSD = 0.6 Å) – green. **c** Single point mutants of HexaPro had increased or decreased binding to 3A3 relative to HexaPro, indicating residues important for 3A3 binding. Each mutant was tested in duplicate in two to five independent ELISA assays. Significance of the difference relative to unmutated HexaPro was determined by ANOVA with post-hoc Tukey-Kramer test with α=0.05 (*) and α=0.01 (**). **d** Location of the mutations that altered binding to 3A3 in the HexaPro spike (6XKL) monomer (left) and **e** in the context of full spike (top down of the S2 portion of the trimer, top; side view of the full spike trimer, bottom). All epitope residues (980–1006) are shown in space-fill, with those not mutated in grey and those mutated without impact on 3A3 binding in black. Locations of mutations that improved binding are displayed in orange and mutations that reduced binding are shown in teal.

### S2 opening provides access to the 3A3 epitope

The epitope mapping and point mutagenesis data indicate that 3A3 binds near the trimer interface of S2, which is poorly accessible in published spike ectodomain structures. Costello/ Shoemaker *et al* have shown that spike undergoes reversible protomer opening in solution to expose the S2 core and the 3A3 epitope^16^. They performed an independent HDX experiment under conditions that favor the open conformation of the trimer and showed that 3A3 does in fact recognize the open state. Consistent with this, 3A3 did not bind a SARS-CoV-2 HexaPro spike that was locked into the closed conformation by engineered disulfide bonds^24^ (Fig. 5a, bottom) although this constrained spike was recognized by the control 2-4 antibody that binds the RBDs in the down state (Fig. 5a, top).

**Figure 5.**
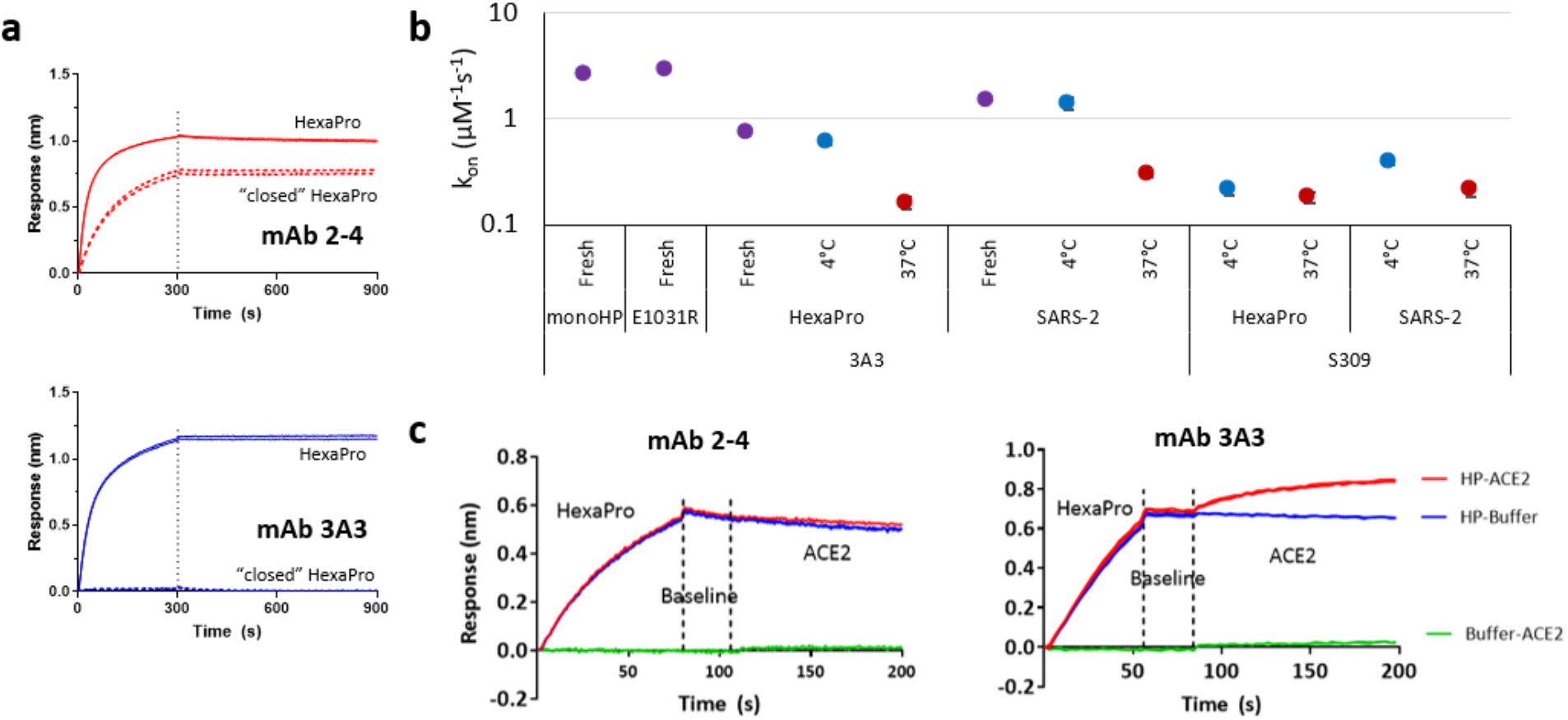
The 3A3 epitope is accessible only when the spike is open. **a** By BLI, the control antibody 2-4^54^ (red lines) bound both SARS-CoV-2 HexaPro (HexaPro, solid) and HexaPro locked into the “closed” conformation (dashed). 3A3 (blue lines) was able to capture HexaPro (solid), but not “closed” HexaPro (dashed). Vertical dashed lines indicate start of the dissociation phase. **b** Antibody 3A3 or S309 was coupled to anti-Fc BLI sensors and allowed to bind monomeric HexaPro (monoHP), E1031R HexaPro (E1031R), HexaPro, or SARS-CoV-2 S2P (SARS-2) spike protein at several concentrations. The proteins were freshly thawed from −80 °C (fresh; purple circles), incubated at 4 °C for one day for SARS-2 or 1 week for HexaPro (4°C; blue circles), or incubated at 37 °C for 1 day (37 °C; red circles). The on-rate of binding (k_on_)was calculated from measurements at seven concentrations in two independent experiments and error is the range of these measurements. **c** Antibody 3A3 or 2-4 were coupled to anti-Fc BLI sensors and allowed to capture HexaPro or nothing (buffer), then dipped into buffer (baseline), and finally dipped into ACE2-Fc (ACE2) or nothing (buffer). In all cases, data were double-reference subtracted.

Analysis of 3A3 binding to various SARS-CoV-2 and SARS-CoV-2 HexaPro spikes with identical 3A3 epitope sequences resulted in inconsistent on-rates and K_d_ values by BLI, likely reflecting epitope accessibility as opposed to the affinity of the epitope/paratope interaction (Fig. 5b, Supplementary Fig. 16). Preincubating spike at 4°C for over one week or 37°C for 24 hours biases spike into the S2-open versus S2-closed conformation, respectively, and correspondingly alters the fraction of spikes with accessible epitope and the apparent 3A3 binding kinetics. The SARS-CoV-2 spike is slightly unstable at 37°C, which alters association of the control antibody S309^22^ for spike by ~2-fold (BLI data; Fig. 5b) and EC_50_ by ~3-fold (ELISA data; Supplementary Fig. 17) relative to spike incubated at 4°C. However, 3A3-SARS-CoV-2 spike binding was markedly reduced after 37°C incubation (~5.5-fold reduced on-rate constant by BLI and ~30-fold increase in ELISA EC_50_). This was also evident in 3A3 binding to SARS-CoV-2 HexaPro after 4°C or 37°C incubation (Fig. 5b). In general, SARS-CoV-2 spike had faster 3A3 on-rate kinetics than SARS-CoV-2 HexaPro, perhaps due to the bias toward the S2-open state in SARS-CoV-2 spike relative to SARS-CoV-2 HexaPro^16^ (Supplementary Fig. 17).

Two additional SARS-CoV-2 HexaPro variants were produced to confirm binding to the S2-open spike state. The E1031R substitution was introduced into SARS-CoV-2 HexaPro, which is outside of the 3A3 epitope but disrupts an electrostatic interaction between E1031 and R1039 on adjacent protomers deep in the S2 base. When evaluated by HDX, this substitution promoted the formation of the S2-open state relative to unmodified HexaPro (Supplementary Fig. 18a). To generate a monomeric HexaPro (monoHP) in which the 3A3 epitope is always exposed, a protease cleavage site was introduced between the C-terminus of the SARS-CoV-2 HexaPro spike and the foldon domain. After 5 days of 4°C incubation, the foldon trimerization motif was cleaved and monoHP isolated by SEC (Supplementary Fig. 18b). When either E1031R or monoHP was assessed for 3A3 binding by BLI, the on-rate was faster than that of fresh, 4°C stored, or 37°C stored unmodified SARS-CoV-2 HexaPro, likely due to improved epitope accessibility (Fig. 5b).

Although ACE2 can bind spike in both the S2-open and S2-closed spike states due to rapid sampling of the RBD up/ down positions, SARS-CoV-2 HexaPro spike shifts toward the S2-open state upon ACE2 binding^16^. It is possible that in the context of unmodified spike on the viral envelope, as in the mammalian cell fusion and pseudovirus neutralization assays (Fig. 2), 3A3 primarily binds when the spike attaches to ACE2. BLI was used to confirm that spike can simultaneously bind ACE2 and 3A3. Immobilized 3A3 captured SARS-CoV-2 HexaPro and then soluble ACE2 (Fig. 5c, right), while mAb 2-4, which binds the RBD down state across protomers, could bind SARS-CoV-2 HexaPro but the complex could not simultaneously bind ACE2 (Fig. 5c, left).

## DISCUSSION

The regular emergence of pathogenic coronaviruses over the past two decades motivated our efforts to isolate cross-reactive coronavirus spike antibodies. In all highly pathogenic coronaviruses, the spike protein is responsible for targeting host cells via the S1 domain, which has little sequence similarity across coronaviruses (41–87%; Supplementary Fig. 1). In contrast, the S2 domain mediates fusion of the viral envelope and target cell membrane through a complex conformational change^2^, with a correspondingly low tolerance for sequence variation (63–98% similarity; Supplementary Fig. 1). Moreover, neutralizing sera from individuals never exposed to SARS-CoV-2 is common in young people and exclusively bind the S2 domain^25^. Accordingly, S2 is an attractive target for pan-coronavirus antibody therapy and vaccination. With this in mind, we isolated and characterized 3A3, a neutralizing and highly cross-reactive antibody binding S2 from a MERS-CoV S2 immune phage library.

Antibody 3A3 binds a highly conserved conformational epitope near the S2 hinge and binds the isolated SARS-CoV-2 HexaPro S2 domain with a low-nanomolar K_d_ by SPR (Table 1 and Supplementary Fig. 4). It exhibits similar binding to the full ectodomain of SARS-CoV-2 HexaPro, SARS-CoV, SARS-CoV-2, and MERS-CoV spikes in ELISA and BLI assays (K_d_ values between 2.5 and 23 nM; Fig. 1, Table 1 and Supplementary Figs. 4 and 5), with minimal binding to the less pathogenic HKU1 spike (Fig. 1a). In addition, 3A3 neutralizes SARS-CoV and SARS-CoV-2 spikes in multiple *in vitro* assays (Fig. 2), with greater potency for the more transmissible D614G and B.1.1.7 variants. The 3A3 binding site was identified by HDX as a cryptic epitope at the apex of S2 (residues 986-1006; Fig. 3). This epitope is conformational, as 3A3 is unable to bind spike stressed by heat and freeze-thaw treatments (Fig. 1). Mutational analysis identified two hot spot residues (D985 and E988) which flank the 2P stabilizing mutations at the “jackknife hinge” between HR1 and the central helix (Fig. 4b-e). This hairpin converts into a long, straight helix during viral fusion, suggesting that 3A3 neutralizes by preventing hairpin extension.

Recent work revealed that 3A3 can only bind SARS-CoV-2 spike when S2 opens to expose the trimer interfaces^16^, suggesting this state is required for 3A3 protection in neutralization assays. Analysis of spike pre-fusion structures shows the 3A3 epitope to be completely obscured when the RBDs are in the three-down or “closed” conformation but is partially exposed when the RBDs become uncoupled from neighboring NTDs^17,21^ and fully exposed when S2 opens^16^. Our biochemical data show that 3A3 does not bind spike locked in the closed, three RBD “down” state (Fig. 5a) but more rapidly associates with multiple SARS-CoV-2 HexaPro variants having exposed S2 interfaces or when spike is treated to favor the S2-open state (Fig. 5b). D614G and B.1.1.7 pseudotyped virus are more readily neutralized by 3A3 than SARS-CoV-2 spike pseudovirus, suggesting that the S2-open versus S2-closed ratio and epitope accessibility are altered in these more infectious variants^26,27^ (Fig. 2c). Conversely, 3A3 only weakly binds the HKU1 spike, which contains three non-conservative changes within the 3A3 epitope (Fig. 4a). None of these seem critical for 3A3 binding (Fig. 4c), suggesting the loss of binding is due to reduced epitope accessibility. For SARS-CoV-2 HexaPro spike, ACE2 binding biases the spike towards the S2-open state^16^, allowing 3A3 to bind the ACE2-spike complex (Fig. 5c), and suggesting that ACE2 engagement may expose the 3A3 epitope in all variants.

Despite success in identifying neutralizing S1 epitopes on SARS-CoV-2 that block ACE2 interactions^6,28–33^, even broadly neutralizing RBD antibodies may be susceptible to escape^34–36^ which has motivated interest in the more conserved S2 domain. Over 200 S2 targeting antibodies have been reported^7^, but very few have been described in detail. Chi *et al.* described over twenty SARS-CoV-2 S2 antibodies, several of which are neutralizing but do not discuss their cross-reactive binding, epitopes, or mechanisms of neutralization^37^. In other work, anti-SARS-CoV-2 S2 antibodies were reported that cross-react with SARS-CoV, MERS-CoV, HKU1 and/or OC43 spike proteins^38^. One antibody binding all except OC43 neutralizes SARS-CoV and SARS-CoV-2 pseudovirus by recognizing an epitope near the S2 base, opposite the 3A3 apex epitope. Additional neutralizing antibodies binding spikes from the three highly pathogenic coronaviruses were reported that bind similar regions on the S2 base^10,11^. By contrast, antibody 3A3, which was elicited by mouse immunization with MERS-CoV S2 and selected with SARS-CoV-2 spike, is highly cross-reactive with the MERS-CoV and SARS-CoV spikes and identifies a novel neutralizing epitope. Interestingly, elicitation of antibodies binding this epitope is not limited to immunization with designed immunogens. An antibody repertoire profiling effort recovered Ab127 from a convalescent COVID-19 patientand used HDX to determine that it protects peptides 980–1006 (overlapping the 3A3 epitope) and 1179–1186 and binds isolated S2 with a 471 nM affinity.^31^ While Ab127 did not neutralize, likely due to its poor affinity, this suggests that the 3A3 epitope is immunogenic during infection.

Broadly neutralizing antibodies binding epitopes on the fusogen trimer interface, similar to 3A3, have been reported recently for other viruses. Conformations analogous to the S2-open state were described for the RSV F^39^ and VSV G^40^ fusogens, suggesting many fusogenic proteins open during conformational change to expose interfacial epitopes. In a few cases, antibodies binding epitopes exposed only in this open state have been described. Antibody D5 and derivatives bind a highly conserved epitope on the N-heptad repeat region of the HIV-1 prehairpin-intermediate to neutralize a wide range of strains^41,42^. Similarly, the broadly neutralizing human antibody A20 recognizes a site at the influenza hemagglutinin head trimer interface to disrupt the trimer^43^. Lee *et al* identified protective anti-flu antibodies that bind different epitopes present on monomeric but not trimeric hemagglutinin^44^. Collectively, these data underscore the novelty of the 3A3 epitope and highlight conserved interfacial epitopes as an emerging strategy for development of broadly neutralizing anti-viral antibodies. As we learn more about the 3A3 paratope, it may serve as a starting point for design of antibodies recognizing even more diverse coronavirus spikes at the structurally conserved S2 hinge.

There are several limitations to this work as currently described. A structure showing the atomic details of 3A3 complexed with spike would provide additional insight into the mechanism of binding and neutralization. Unfortunately, structures of antibodies bound to S2 are generally challenging to obtain with just two high-resolution structures of antibody-S2 peptide complexes available^10,11^ and none in complex with the full spike ectodomain. Antibody 3A3 has a ~2.5 nM affinity for purified SARS-CoV-2 spike with two stabilizing prolines, but neutralization IC_50_ values with the unstabilized homolog are >10-fold higher. Unstabilized spike may less frequently sample the S2-open state, and the stabilizing prolines may rigidify the hairpin epitope or subtly alter presentation of hot spot residues D985 and E988. Affinity maturation of 3A3 to bind SARS-CoV-2 spike without the K986P and V987P changes may yield a more potent broadly neutralizing antibody and help overcome accessibility limitations in spike variants not favoring the S2 open state.

Here we report an antibody binding a neutralizing S2 epitope that is conserved across all highly pathogenic coronavirus strains. HDX mapping and epitope mutagenesis identified the epitope in the hinge at the S2 apex, suggesting 3A3 inhibits the pre- to post-fusion spike transition. The 3A3 antibody binds spike when the S2 domain trimer opens, a state which exists in equilibrium with the S2-closed prefusion state in stabilized spikes. Multiple deployed vaccines utilize versions of the 2P stabilized SARS-CoV-2 spike, and antibodies binding 3A3-like epitopes in the S2 core may be elicited during immunization. The relevance of these antibodies in both convalescent and immunized individuals and for future protection from novel coronaviruses is important to understand. Accordingly, future work will focus on affinity maturation of 3A3 and high-resolution structural analysis to determine the precise antibody footprint and neutralizing mechanism.

## MATERIALS AND METHODS

### Spike Expression

Soluble coronavirus spikes and spike variants were expressed and purified as previously described ^17,21^. SARS-CoV-2^17^, SARS-CoV^45^, SARS-CoV-2 HexaPro^21^ spikes and epitope variants, HexaPro E1031R, and monoHP were expressed in ExpiCHO cells (ThermoFisher Scientific). The monoHP variant was produced by adding a short linker and a HRV3C cleavage site between the C-terminus of SARS-CoV-2 HexaPro and the T4 fibritin domain. The purified protein was incubated at 4°C for ~5 days and incubated overnight with human rhinovirus 3C (HRV3C) protease. The monoHP spike was purified by SEC from remaining trimer, HRV3C protease and tags. MERS-CoV^46^, HKU1^46^, and the SARS-CoV-2 variants HexaPro S2 (residues 697–1208 of the SARS-CoV-2 spike with an artificial signal peptide, proline substitutions at positions 817, 892, 899, 942, 986 and 987 and a C-terminal T4 fibritin domain, HRV3C cleavage site, 8xHisTag and TwinStrepTag), HexaPro RBD-locked-down (HexaPro with S383C-D985C substitutions), and aglycosylated HexaPro (HexaPro treated with Endo H overnight at 4°C leaving only one N-acetylglucosamine attached to N-glycosylation site) as well as MERS-CoV S2-only (residues 763–1291 of MERS-2P with 8 additional stabilizing substitutions), MERS-CoV S2-apex-less (MERS-CoV S2-only construct with residues 811–824 replaced with GGSGGS and residues 1042–1073 replaced with a flexible linker) were expressed in Freestyle 293-F cells (ThermoFisher Scientific).

### Murine immunization

Three BALB/c mice were immunized subcutaneously with 5 μg pre-fusion stabilized MERS-CoV S2 and 20 μg of ODN1826 + 100 μl of 2X Sigma Adjuvant System (SAS; Sigma) containing monophosphoryl lipid A and trehalose dimycolate in squalene oil. Four weeks later, the mice were boosted with the same dose of the same mixture. Three weeks after boosting, the mice were sacrificed and spleens were collected in RNALater (ThermoFisher). The University approved mouse protocols of Texas at Austin IACUC (AUP-2018-00092).

### Phage display antibody library construction

RNA was isolated from the aqueous phase of homogenized spleens mixed with 1-bromo-3-chloropropane and purified with the PureLink RNA kit (Invitrogen) separately. The Superscript IV kit (Invitrogen) was used to synthesize cDNA. The V_H_ and V_L_ sequences from each immunized mouse were amplified with mouse-specific primers described by Krebber *et al*^18^. Maintaining separate reactions for each mouse, the V_L_ and V_H_ regions were joined by overlap extension PCR for each immunized mouse spleen to generate VL-linker-VH fragments (scFv) which the linker region encodes the amino acids (Gly_4_Ser)_4_ and *SfiI* sites flanked the scFv sequence. The scFv PCR products were pooled and cloned into pMopac24^19^ via *SfiI* cut sites to encode an scFv with a c-terminal myc tag fused to the M13 phage pIII protein. This library was then transformed to XL1-Blue (Agilent Technologies) *E. coli.* The total number of transformants was 3.1×10^8^ with <0.01% background based on plating.

### Phage display and panning

The *E.coli* containing the library were expanded in growth media (2×YT with 1% glucose, 200 μg/mL ampicillin, 10 μg/mL tetracycline) at 37 °C to an OD_600_ of 0.5, then infected with 1 × 10^11^ pfu/mL M13K07 helper phage (NEB) and induced with 1 mM isopropyl β-d-1-thiogalactopyranoside. After two hours of shaking at room temperature, 12.5 μg/mL of kanamycin was added for phage expression overnight. Phage were precipitated in 20% PEG-8000 in 2.5 M NaCl, titered by infection of XL1-Blue and plating, and used for Round 1 panning. This process was repeated for each round of panning, starting from overnight growth of the output phage from each round.

Four rounds of panning were used to isolate scFvs binding both MERS-CoV S2 and SARS-CoV-2 spike using the following solutions coated on high binding plates: 2 μg/mL anti-c-myc tag antibody (Invitrogen) to eliminate phage expressing no or truncated scFv (Round 1), 2 μg/mL MERS-CoV S2 (Round 2), 2 μg/mL SARS-CoV-2 spike (Round 3), and 0.4 μg/mL SARS-CoV-2 spike (Round 4). In each round of panning, the plates were blocked with 5% non-fat milk in phosphate-buffered saline (PBS) with 0.05% Tween-20 (PBS-T), and phage were preincubated with 5% non-fat milk in PBS-T for 30 minutes before incubation on the plate for 1.5 hours at room temperature. After thorough washing with PBS-T, output phage was eluted using 0.1 M HCl at pH 2.2, neutralized with ~1:20 2 M Tris base, and allowed to infect XL1-Blue cells overnight amplification.

Random clones isolated after Round 3 and Round 4 of panning were sequenced and unique clones were tested by monoclonal phage enzyme-linked immunosorbent assay (ELISA) on plates coated with SARS-CoV-2 spike or RSV F foldon at 2 μg/mL in PBS. Briefly, plates were coated overnight at 4°C, washed with PBS-T, then blocked with PBS-T and 5% milk. Phage were allowed to bind for one hour at room temperature, thoroughly washed with PBS-T, then incubated with 1:2000 anti-M13 pVIII-HRP (GE Healthcare) in PBS-T 5% milk for another hour. After washing, the plate was developed with the TMB Substrate Kit (Thermo Scientific), quenched with an equal volume of 1 M HCl and evaluated by absorbance at 450 nm (Supplementary Fig. 2).

### Antibody expression, purification, and quality control

Full-length antibody versions of 3A3, 4A5, 4H2, and 3E11 were cloned as previously described^47^ as mouse variable region-human IgG1 constant region chimeras. Antibodies were expressed in ExpiCHO (ThermoFisher Scientific) cells according to the high titer protocol provided and purified on a Protein A HiTrap column (GE Healthcare) with the ACTA Pure FPLC system (GE Healthcare), and buffer exchanged to PBS. Each purified antibody was analyzed by SDS-PAGE (3 μg antibody per well) under reducing and non-reducing conditions (Supplementary Fig. 3a) and by analytical SEC on a Superdex S200 column (GE Healthcare) (Supplementary Fig. 3b).

Mouse Fab fragments of each sequence were generated by cloning the V_H_ and V_L_ regions upstream of heavy chain constant regions with a HRV3C protease site in the hinge^46^ and a mouse kappa chain, respectively. After expression, protein A purified protein was digested with HRV3C protease, and the flow-through from a protein A HiTrap column was collected. Excess HRV3C protease was removed by incubation with Ni Sepharose 6Fast Flow beads (GE Healthcare). Fully murine antibodies were produced by cloning the VH regions into a mouse IgG2 expression cassette in the pAbVec background, co-transfected with the appropriate mouse IgK plasmid ^48^, and purified as described above.

According to the kit instructions, the thermal unfolding temperatures of the chimeric antibodies (0.3 mg/mL) were assessed in triplicate using the Protein Thermal Shift Dye Kit (ThermoFisher Scientific). Continuous fluorescence measurements (λ_ex_ = 580 nm, λ_em_ = 623 nm) were performed using a ThermoFisher ViiA 7 Real-Time PCR System, with a temperature ramp rate of 0.05°C/sec increasing from 25°C to 99°C (Supplementary Fig. 3c and d).

### ELISA evaluation of antibody cross-reactivity and binding to stressed spike

ELISAs were performed as described above throughout the work. For testing each antibody’s specificity, plates were coated with 1 μg/mL of purified spike proteins (SARS-CoV, SARS-CoV-2, SARS-CoV-2 HexaPro, MERS-CoV, HKU1, and RSV F foldon) in PBS. Duplicate serial dilutions of each full-length antibody were allowed to bind each coat, and the secondary antibody solution was a 1:1200 dilution of goat-anti-human IgG Fc-HRP (SouthernBiotech). ELISA curves were fit to a 4-parameter logistic curve (Fig. 1a and Supplementary Fig. 5a and b).

To stress the spike proteins, fresh aliquots of SARS-CoV-2 and SARS-CoV-2 HexaPro spikes were thawed and split. One half of the aliquot was stressed by incubation at −20°C for 5 min, then 50°C for 2 min for a total of three temperature cycles. The freshly thawed and stressed spikes were serially diluted and captured on ELISA plates coated with each full-length antibody at 1 μg/mL or nothing (no coat). The ELISA was carried out as above with 3% w/v BSA in place of milk in the diluent and blocking buffer and Strep-Tactin-HRP (IBA) as the secondary reagent (Fig. 1c and d, and Supplementary Fig. 5d). For each fresh and stressed spike, 8 μg was analyzed by SDS-PAGE under non-reducing conditions (Supplementary Fig. 6).

### Western blot of antibody binding to coronavirus spike proteins

Purified coronavirus spike proteins (SARS-CoV-2 HexaPro, SARS-CoV-2, MERS-CoV, and HKU1) were reduced and boiled, and 50 ng of each was subjected to SDS-PAGE and transfer to PVDF membranes in quadruplicate. After blocking with PBS-T with 5% milk, the membranes were probed with 0.2 μg/mL 3A3, 1 μg/mL 4A5, 1 μg/mL 4H2 or 0.2 μg/mL 3E11 for 1 hour at room temperature. After washing with PBST, the membranes were incubated with 1:4000 goat anti-human IgG Fc-HRP for 45 minutes at room temperature, then developed with the SuperSignal West Pico Chemiluminescent Substrate (Thermo Scientific) and imaged (Fig. 1b and Supplementary Fig. 5c).

### Surface plasmon resonance (SPR) and biolayer interferometry (BLI) measurements

SPR was used to determine the binding kinetics and affinity of the 3A3 Fab and HexaPro S2 interaction. An anti-StrepTagII Fab was covalently coupled to a CM5 sensor chip, which was then used to capture purified HexaPro S2 by the c-terminal twin StrepTag to ~80 response units (RU) in each cycle using a Biacore X100 (GE Healthcare). The binding surface was regenerated between cycles using 0.1% SDS followed by 10 mM Glycine at pH 2. The 3A3 Fab was serially diluted from 12.5 nM to 1.56 nM and injected over the blank reference flow cell and then HexaPro S2-coated flow cell in HBS-P+ buffer. Buffer was also injected through both flow cells as a reference. The data were double-reference subtracted and fit to a 1:1 binding model using BIAevaluation software.

To determine the affinity of 3A3 Fab by BLI, anti-human IgG Fc (AHC) (ForteBio) sensors were coated with the anti-foldon antibody identified in this work (3E11) at 10nM in the kinetics buffer (0.01% BSA and 0.002% Tween-20 in PBS) to a response of 0.6 nm. MAb-coated sensors were then incubated with HexaPro S2 at 60 nM to a response of 0.6 nm. Association of 3A3 Fab was recorded for 5 minutes in kinetics buffer, starting at 100 nM followed by 1:2 dilutions. The dissociation was recorded for 10 minutes in the kinetics buffer. K_d_ values were obtained using a 1:1 global fit model using the Octet instrument software. 3A3 Fab kinetics measurement was repeated once (Supplementary Fig. 4b and Table 1).

To determine the apparent K_d_ values of IgGs, anti-Human IgG Fc sensors were loaded with mAbs in the kinetics buffer at 10 nM to a response of 0.6 nm. Association curves were recorded for 5 minutes by incubating the sensors in different concentrations of spike or spike S2 domains, starting from 100 nM and serially diluted at 1:2. The dissociation step was recorded for 10 minutes in the kinetics buffer. Steady-state K_d_ values were determined using the response values obtained after five minutes of association using the Octet analysis software (Table 1 and Supplementary Figs. 4c and d). To evaluate spikes incubated at different temperatures, the spikes were stored for ~1–3 weeks at 4°C or 24 hours at 37°C, then diluted into room temperature buffer immediately before measurements.

To compare 3A3 and mAb 2-4 binding to HexaPro and “Down” HexaPro, anti-Human IgG Fc sensors were loaded with 3A3 or mAb 2-4 mAb in the kinetics buffer at 10 nM to a response of 0.6 nm. After a baseline step, the sensors were incubated with either HexaPro or “Down” HexaPro, both at 60 nM for five minutes. Dissociation step was recorded for 10 minutes in the kinetics buffer (Fig. 5a).

To evaluate ACE2 binding to HexaPro captured by 3A3, Anti-Human Fc Sensors were used to pick up 3A3 (10 nM) to a response of 0.6 nm. Then mAb coated tips were dipped into wells containing HexaPro (50 nM) to a response of 0.6 nm and then dipped into wells containing ACE2 (50 nM), irrelevant murine mAb (50 nM), or buffer. Association of mu3A3/irrelevant mAb was measured for 5 minutes and dissociation for 10 minutes. (Fig. 5c).

Octet Red96 (ForteBio) instrument was used. Between every loading step, sensors were washed with kinetics buffer for 30 seconds. Before use, sensors were hydrated in the kinetics buffer for 10 minutes. After each assay, the sensors were regenerated using 10 mM Glycine, pH 1.5.

### Confocal cell fusion assay

On day 0, the CHO-T cells (Acyte Biotech) were transfected with either pPyEGFP ^49^, 1:4 pWT-SARS-CoV-2-spike:pPyEGFP, and 1:4 pD614G-SARS-CoV-2-spike:pPyEGFP using Lipofectamine 2000 (Life Technologies), and media was replaced on day 1. On day 2 after transfection, HEK-293T-hACE2 cells (BEI, NR-52511), which stably expresses human ACE2, were stained with 1 μM CellTrace Far Red dye (Invitrogen, Ex/Em: 630/661 nm) in PBS for 20 minutes at room temperature, then quenched with DMEM with 10% heat-inactivated FBS for 5 minutes, and resuspended in fresh media. CHO-T cells expressing EGFP or EGFP and surface spike were preincubated with the antibody for one hour at 37°C, then mixed with HEK-hACE2 cells at a ratio of 5:1 in 24-well plates with a coverslip on the bottom of each well. On day 3, after 20 hours of coincubation, the coverslip with bound cells was washed once with PBS and fixed with 4% paraformaldehyde for 20 min at room temperature, washed again, and mounted on slides with DAPI-fluoromount-G (Southern Biotech). Images were collected with Zeiss LSM 710 confocal microscope (Carl Zeiss, Inc) and processed using ImageJ software (http://rsbweb.nih.gov/ij) (Fig. 2 and Supplementary Fig. 7).

The cell fusion level was determined by two different statistical analysis methods. The first statistical analysis was based on the percentage of HEK-ACE2 pixels (red) colocalizing with spike expressing CHO pixels (green), which was determined by the following equation within the JACoP plugin for ImageJ ^50^:

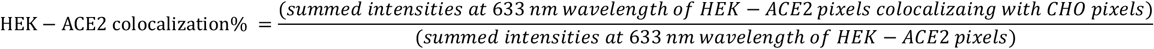

The colocalization percentage for each independent image was determined using the Manders’ coefficient. The second statistical analysis was based on the average HEK-ACE2 cell size after the coincubation with CHO cells using the ImageJ software. The image of HEK-ACE2 (red fluorescence color at 633 nm wavelength) was converted into 16-bit in greyscale and adjusted the threshold to highlight the cell structure. The average cell size was automatically counted with the “Analyze Particles” tab with a size threshold (50-infinity) to exclude the background noise. The cell on the edge was excluded. The statistical significance of either HEK-ACE2 colocalization percentage or average cell size between different conditions was calculated with ANOVA using GraphPad Prism 7 (GraphPad Software). Values represent the mean and standard deviation of at least 160 cells.

### Lentiviral plasmids

Plasmids required for lentiviral production were obtained from BEI Resources. Plasmids expressing the HIV virion under the CMV promotor (HDM-Hgpm2, pRC-CMV-Rev1b, and HDM-tat1b) were provided under the following catalog numbers NR-52516, NR-52519, and NR-52518, respectively^51^. Plasmids for lentiviral backbone expressing a luciferase reporter under the CMV promotor followed by an IRES and ZsGreen (pHAGE-CMV-Luc2-IRES-ZsGreen-W) or human ACE2 gene (GenBank ID NM_021804) under an EF1a promoter (pHAGE2-EF1aInt-ACE2-WT) were provided as NR-52520 and NR52516, respectively^51^. The envelop vector expressing a codon-optimized wild-type SARS-CoV-2 spike protein (Genbank ID NC_045512) under a CMV promoter was obtained from BEI resources (HDM-IDTSpike-fixK, NR-52514)^51^, while the plasmid expressing VSV G (vesicular stomatitis virus glycoprotein) was purchased from Cell Biolabs (pCMV-VSV-G, Part No. RV-110). The HDM-IDTSpike-fixK plasmid was employed as a template for site-directed mutagenesis to generate the expression plasmid for the D614G mutant of SARS-CoV-2 spike protein. SARS-CoV and B.1.1.7 spikes were cloned into the same HDM-IDTSpike-fixK plasmid for pseudovirus production.

### Generation of HEK293T-ACE2 target cells, stably expressing human ACE2

A lentiviral vector (pHAGE2-EF1aInt-ACE2-WT) expressing human ACE2 under an EF1a promoter was used to transduce HEK293T cells. Clonal selection depended on the susceptibility to infection by the pseudotyped lentiviral particles; selected clones were validated using western blotting.

### SARS-CoV-2 spike-mediated pseudovirus entry assay

HIV particles pseudotyped with wild-type, D614G or B.1.1.7 variants of SARS-CoV-2 spike and SARS-CoV spike were generated in HEK293T cells. A detailed protocol for generating these particles was reported by Crawford et al. ^51^. HEK293T cells were co-transfected with plasmids for (1) HIV virion-formation proteins (HDM-Hgpm2, pRC-CMV-Rev1b, and HDM-tat1b; (2) lentiviral backbone expressing luciferase reporter (pHAGE-CMV-Luc2-IRES-ZsGreen-W), and (3) a plasmid encoding one of the envelope proteins (wild-type SARS-CoV-2, D614G mutant, B.1.1.7 variant, SARS-CoV or VSV G). 72 hours post-transfection, media containing the pseudovirus particles were collected, filtered, fractionated, and stored at −80 °C. The particles were used directly in cell entry experiments or after pre-incubation with each antibody for one hour at room temperature. After 60–72 hours, a total number of cells per well were estimated using an lncuCyte® ZOOM equipment with a ×10 objective. Then cells were treated with the Bright-Glo Luciferase Assay reagent (Promega, E2610) to detect a luciferase signal (relative luciferase units or RLU) following the manufacturer’s protocol. The percentage of entry was estimated as the ratio of the relative luciferase units recorded in the presence and absence of the tested antibody and a half-maximal inhibitory concentrations (IC_50_) calculated using a 3-parameter logistic regression equation (GraphPad Prism v9.0) (Fig. 2).

### Flow cytometry

On day 0, Expi-293 cells (ThermoFisher) were mock transfected or transfected with pWT-SARS-2-spike or pD614G-SARS-2-spike described for pseudovirus assays above. On day 2, 30 nM full-length 3A3 or CR3022 was added to ~5×10^5^ transfected cells for 1 hour on ice. All cells were collected, washed with PBS with 1% FBS, then incubated with 1:100 goat-anti-human Fc-AF647 for one hour on ice. Cells were washed again, then scanned for AF647 (640 nm excitation, 670/30 bandpass emission) fluorescence on a BD Fortessa flow cytometer and analyzed with FlowJo (Supplementary Fig. 9).

### Hydrogen-Deuterium Exchange Mass Spectrometry

Hydrogen-deuterium exchange was performed on 0.50 μM SARS-CoV-2 HexaPro spike protein alone (Supplementary Fig. 11) or in the presence of 0.55 μM 3A3 IgG or Fab (Fig. 3 and Supplementary Fig. 12), 0.75 μM 4A5, or 0.75 μM 4H2 (Supplementary Fig. 13). Complexes were incubated for 10 min at 25°C before exchange in 90% deuterium and 20 mM Tris pH 8.0, 200 mM NaCl. The exchange was quenched after 10^1^, 10^2^, 10^3^ and 10^4^ s by mixing samples 1:1 with cooled 0.2% (v/v) Formic acid, 200 mM TCEP, 8 M Urea, pH 2.3. Samples were immediately flash-frozen in liquid N_2_ and stored at −80°C.

Samples were thawed and LC-MS performed using a Waters HDX manager and SYNAPT *G2-Si* Q-Tof. Three or four technical replicates of each sample were analyzed in random order. Samples were digested on-line by *Sus scrofa* Pepsin A (Waters Enzymate™ BEH Pepsin column) at 15°C and peptides trapped on a C18 pre-column (Waters ACQUITY UPLC BEH C18 VanGuard pre-column) at 1°C for 3 min at 100 μL/min. Peptides were separated over a C18 column (Waters ACQUITY UPLC BEH C18 column) and eluted with a linear 3–40% (v/v) Acetonitrile gradient for 7 min at 30 uL/min at 1 °C and 0.1% (v/v) Formic Acid as the basic LC buffer.

MS data were acquired using positive ion mode and either HDMS or HDMS^E^. HDMS^E^ mode was used to collect both low (6 V) and high (ramping 22–44 V) energy fragmentation data for peptide identification in water-only samples. HDMS mode was used to collect low energy ion data for all deuterated samples. All samples were acquired in resolution mode. The capillary voltage was set to 2.8 kV for the sample sprayer. Desolvation gas was set to 650 L/hour at 175°C. The source temperature was set to 80°C. Cone and nebulizer gas were flowed at 90 L/hour and 6.5 bar, respectively. The sampling cone and source offset were set to 30 V. Data were acquired at a scan time of 0.4 s with a range of 100–2000 m/z. A mass correction was done using [Glu1]-fibrinopeptide B as a reference mass.

Water-only control samples were processed by Protein Lynx Global Server v.3.0.2 with a ‘minimum fragment ion matches per peptide’ of 3 and allowing methionine oxidation. The low and elevated energy thresholds were 250 and 50 counts, respectively, and the overall intensity threshold was 750 counts. The resulting peptide lists were then used to search data from deuterated samples using DynamX v.3.0. Peptide filters of 0.3 products per amino acid and 1 consecutive product were used. Spectra were manually assessed, and figures were prepared using HD-eXplosion^52^ and PyMOL^53^. The HDX data summary table (Supplementary Table 1) and complete data table (Supplementary Table 2) are included.

### Statistical analyses

The means ± SD were determined for all appropriate data. For the mammalian cell fusion experiments, pseudovirus neutralization experiments and epitope variant analysis, a one-way analysis of variance (ANOVA) with Tukey’s simultaneous test with *P* values was used to determine statistical significance between groups. Welch’s t-test was used to determine the significance of deuterium uptake differences.

## General

The following reagent was contributed by David Veesler for distribution through BEI Resources, NIAID, NIH: Vector pcDNA3.1(-) containing the SARS-Related Coronavirus 2, Wuhan-Hu-1 Spike Glycoprotein Gene, NR-52420 ^55^. The following reagents were obtained through BEI Resources, NIAID, NIH: Human Embryonic Kidney Cells (HEK-293T) Expressing Human Angiotensin-Converting Enzyme 2, HEK-293T-hACE2 Cell Line, NR-52511; SARS-Related Coronavirus 2, Wuhan-Hu-1 Spike-Pseudotyped Lentiviral Kit, NR-52948 (including NR-52514, NR-52516, NR-52517, NR-51518, and NR-52519); and Vector pHAGE2 Containing the ZsGreen Gene, NR-52520.

The authors would like to thank Eduardo A. Padlan, Gregory C. Ippolito, Jason J. Lavinder, and George Delidakis for useful discussions and advice related to this work.

## Author contributions

Conceptualization, Y.H., A.W.N., C.-L.H., A.N.Q., S.R.S., S.M.C., S.M., J.S.M. and J.A.M.; Investigation and visualization, Y.H., A.W.N., C.-L.H., R.S., O.S.O., R.W., T.S.K., L.R.A., A.N.Q., K.C.L., A.L.B., A.M.D., Y.L., A.L., D.A., S.R.S. and S.M.C.; Writing –Original Draft, Y.H., A.W.N, T.S.K., S.D. and J.A.M.; Writing – Reviewing & Editing, Y.H., A.W.N., C.-L.H., R.S., O.S.O., R.W., T.S.K., S.M., K.D., S.D., J.S.M and J.A.M.; Supervision, S.M., K.D., S.D., J.S.M. and J.A.M.

## Data and materials availability

Data and antibody sequences will be submitted to the Submit data to the Immune Epitope Database and Analysis Resource (iedb.org) and the Coronavirus Immunotherapy Consortium (covic.lji.org).

