## Supplemental materials for "Identification of a conserved neutralizing epitope present on spike proteins from all highly pathogenic coronaviruses"

Yimin Huang[1]†, Annalee W. Nguyen[2]†, Ching-Lin Hsieh[1], Rui Silva[1], Oladimeji S. Olaluwoye[4], Rebecca E. Wilen[2], Tamer S. Kaoud[3], Laura R. Azouz[2], Ahlam N. Qerqez[2], Kevin C. Le[2], Amanda L. Bohanon[1], Andrea M. DiVenere[2], Yutong Liu[2], Alison G. Lee[1], Dzifa Amengor[1], Sophie R. Shoemaker[5], Shawn M. Costello[6], Susan Marqusee[5,7], Kevin N. Dalby[3], Sheena D'Arcy[4], Jason S. McLellan[1,8]*, Jennifer A. Maynard[2,8]*

**a** S1 Identity

| | 229E | NL63 | OC43 | HKU1 | MERS | SARS 1 | SARS 2 |
|---|---|---|---|---|---|---|---|
| 229E | 100% | 50% | 21% | 21% | 21% | 20% | 22% |
| NL63 | | 100% | 21% | 20% | 21% | 20% | 21% |
| OC43 | | | 100% | 57% | 24% | 24% | 23% |
| HKU1 | | | | 100% | 24% | 24% | 23% |
| MERS | | | | | 100% | 24% | 21% |
| SARS 1 | | | | | | 100% | 65% |
| SARS 2 | | | | | | | 100% |

**b** S2 Identity

| | 229E | NL63 | OC43 | HKU1 | MERS | SARS 1 | SARS 2 |
|---|---|---|---|---|---|---|---|
| 229E | 100% | 76% | 35% | 35% | 36% | 35% | 35% |
| NL63 | | 100% | 37% | 35% | 36% | 33% | 33% |
| OC43 | | | 100% | 72% | 46% | 42% | 42% |
| HKU1 | | | | 100% | 43% | 40% | 41% |
| MERS | | | | | 100% | 42% | 44% |
| SARS 1 | | | | | | 100% | 90% |
| SARS 2 | | | | | | | 100% |

**c** S1 Similarity

| | 229E | NL63 | OC43 | HKU1 | MERS | SARS 1 | SARS 2 |
|---|---|---|---|---|---|---|---|
| 229E | 100% | 78% | 44% | 47% | 44% | 46% | 41% |
| NL63 | | 100% | 44% | 42% | 44% | 44% | 45% |
| OC43 | | | 100% | 80% | 49% | 49% | 49% |
| HKU1 | | | | 100% | 52% | 51% | 50% |
| MERS | | | | | 100% | 52% | 51% |
| SARS 1 | | | | | | 100% | 87% |
| SARS 2 | | | | | | | 100% |

**d** S2 Similarity

| | 229E | NL63 | OC43 | HKU1 | MERS | SARS 1 | SARS 2 |
|---|---|---|---|---|---|---|---|
| 229E | 100% | 93% | 65% | 66% | 64% | 66% | 67% |
| NL63 | | 100% | 67% | 65% | 63% | 65% | 67% |
| OC43 | | | 100% | 93% | 71% | 71% | 72% |
| HKU1 | | | | 100% | 72% | 72% | 73% |
| MERS | | | | | 100% | 73% | 74% |
| SARS 1 | | | | | | 100% | 98% |
| SARS 2 | | | | | | | 100% |

**Supplementary Figure 1. Sequence conservation is higher in the S2 domain than the S1 domain across coronaviruses that infect humans.** Percent sequence identity and similarity between the spike (**a** and **c**, respectively) S1 subunits and (**b** and **d**, respectively) S2 subunits of the seven coronaviruses known to infect humans was analyzed using LALIGN/PLALIGN local alignment (https://fastademo.bioch.virginia.edu/fasta_www2/fasta_www.cgi?rm=lplalign). The default values of the gap penalties of open = -12 and gap = -2 were used for S2 alignments but reduced to open = -10 and gap = -1 to allow for variable domain lengths in S1. GenBank protein sequence identification numbers (protein_id) used in these alignments were: NP_073551.1 for 229E, AFD64754.1 for NL63, BBA20979.1 for OC43, BBA20986.1 for HKU1, YP_009047204.1 for MERS, AYV99817.1 for SARS-CoV-1, and YP_009724390.1 for SARS-CoV-2.

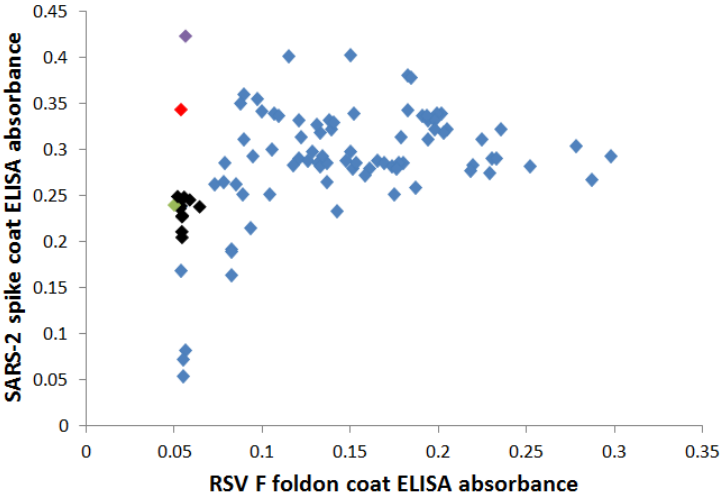


**Supplementary Figure 2. Many cross-reactive scFv-phage target the foldon domain.** The majority of monoclonal phage tested by ELISA on SARS-CoV-2 spike or the unrelated RSV F foldon coated plates had cross-reactive binding, indicating targeting of the shared foldon domain. After round 4 of panning, these data show 3A3 in black, a close relative of 3A3 (two amino acid changes) in green, 4A5 in red, 4H2 in purple, and foldon binders with closely related CDRH3 sequences in blue.

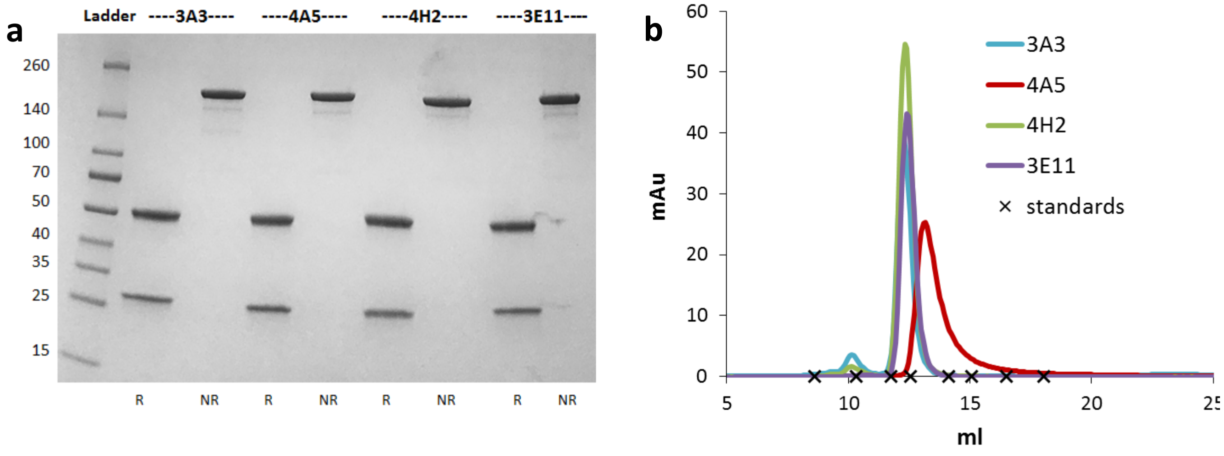


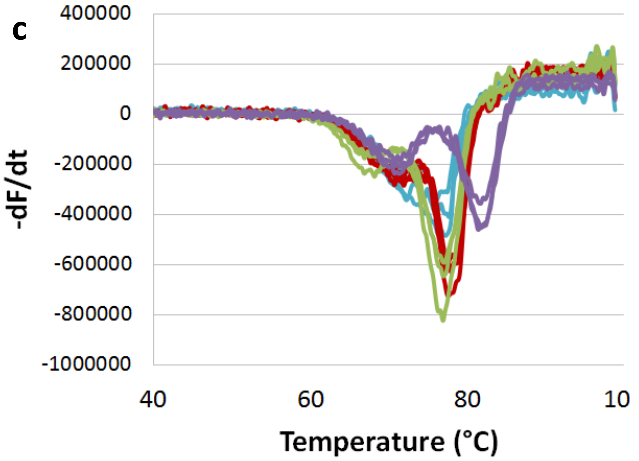


d

| | IgG1 $T_m$ (°C) | |
|---|---|---|
| **antibody** | 1st transition | 2nd transition |
| **3A3** | 72.0 ± 0.3 | 76.9 ± 0.8 |
| **4A5** | 71.4 ± 0.7 | 77.6 ±0.3 |
| **4H2** | 67.5 ± 0.6 | 76.9 ±0.1 |
| **3E11** | 70.9 ± 0.3 | 81.6 ±0.2 |

**Supplementary Figure 3. The purified 3A3, 4A5, 4H2, and 3E11 full-length antibodies are pure and stable. a** SDS-PAGE was used to evaluate 3 µg of each antibody in either reduced (R) or non-reduced (NR) states. Molecular weights of each ladder band in kDa are indicated on the left. **b** 100 µg of each antibody was evaluated by analytical SEC on an S200 column. Standards (marked with an x) are: thyroglobulin (669 kDa, 8.609 ml); ferritin (440 kDa, 10.316 ml); beta-amylase (200 kDa, 11.743 ml); aldolase (158 kDa, 12.574 ml); conalbumin (75 kDa, 14.131 ml); ovalbumin (44 kDa, 15.079 ml); carbonic anhydrase (29 kDa, 16.484 ml); cytochrome c (12.4 kDa, 18.046 ml). **c** Thermal unfolding of 3A3 (blue), 4A5 (red), 4H2 (green), and 3E11 (purple) show profiles typical of antibodies. **d** Average and standard deviations (n=3) of the first and second transition temperatures were identified from minima in -dF/dt during thermal unfolding.

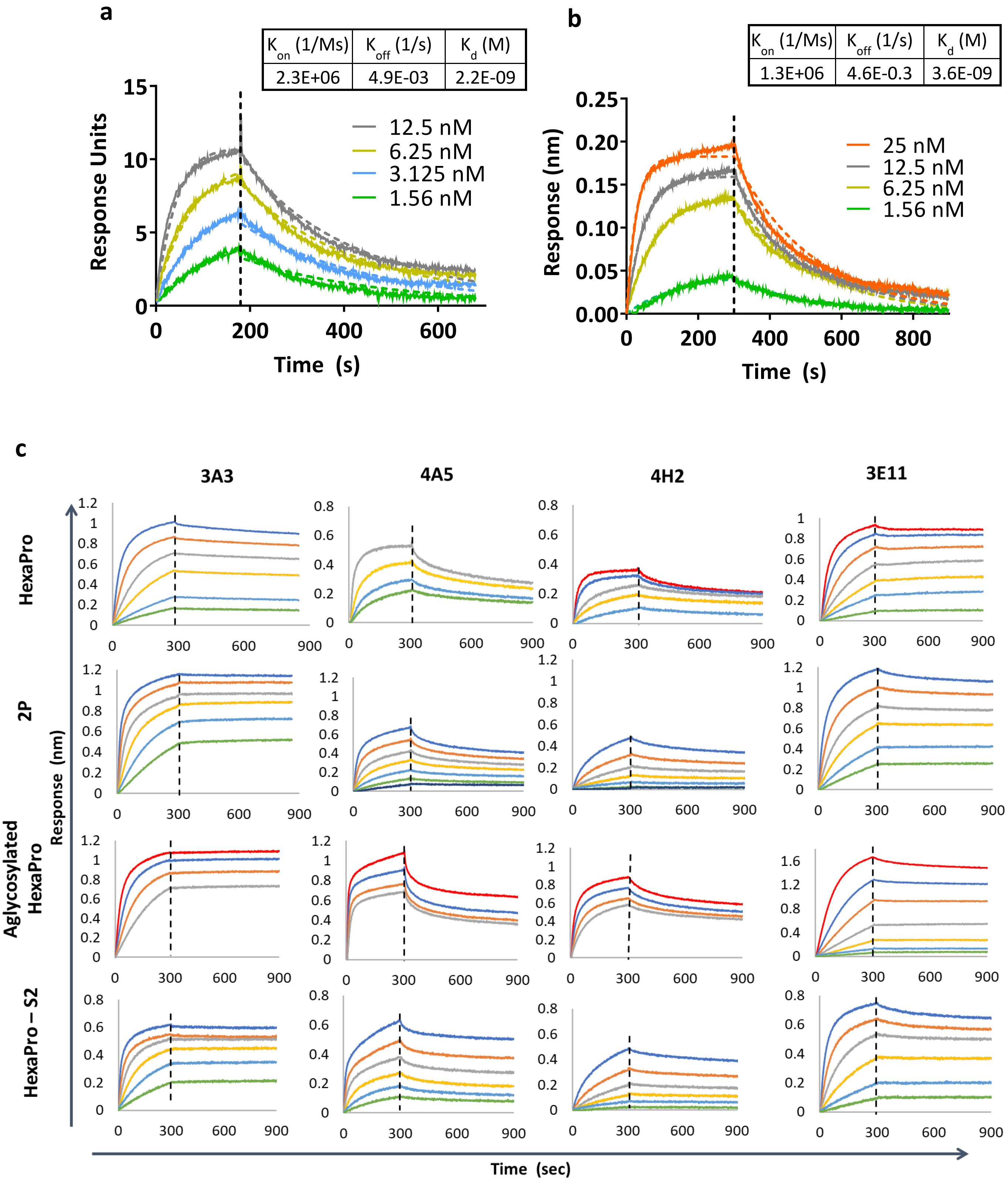
a
K on (1/Ms) | K off (1/s) | K d (M)
2.3E+06 | 4.9E-03 | 2.2E-09
Response Units
Time (s)
12.5 nM
6.25 nM
3.125 nM
1.56 nM
b
K on (1/Ms) | K off (1/s) | K d (M)
1.3E+06 | 4.6E-0.3 | 3.6E-09
Response (nm)
Time (s)
25 nM
12.5 nM
6.25 nM
1.56 nM
c
3A3
4A5
4H2
3E11
HexaPro
2P
Aglycosylated HexaPro
HexaPro – S2
Response (nm)
Time (sec)

d

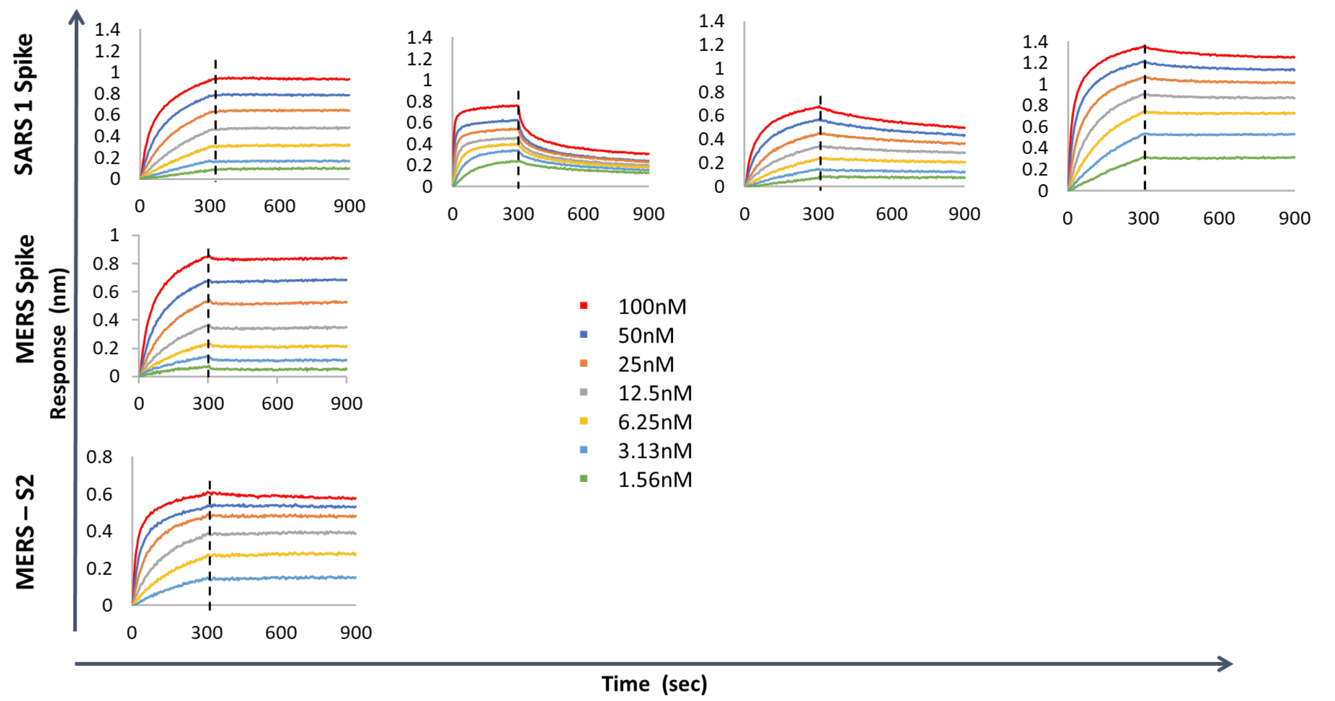


**Supplementary Figure 4. SPR and BLI analysis of 3A3 Fab. 3A3, 4A5, 4H2 and 3E11 mAbs measured low to mid nanomolar affinity ($K_d$) for binding to SARS-CoV-2 spike variants.** Binding of 3A3 Fab to HexaPro S2 measured by SPR **a** analysis and BLI **b**. Equilibrium affinity of immobilized full-length IgGs on anti-Fc sensors capturing the indicated **c** SARS-CoV-2 spike or S2 domain and **d** SARS-CoV or MERS-CoV spike. Measurements were performed by dipping full-length antibody-coated sensors into serial dilutions of the different spike variants (100 to 1.56 nM) followed by a dissociation step in the buffer. Vertical lines indicate the start of the dissociation phase; data shown in solid lines with fits shown in dashed lines.

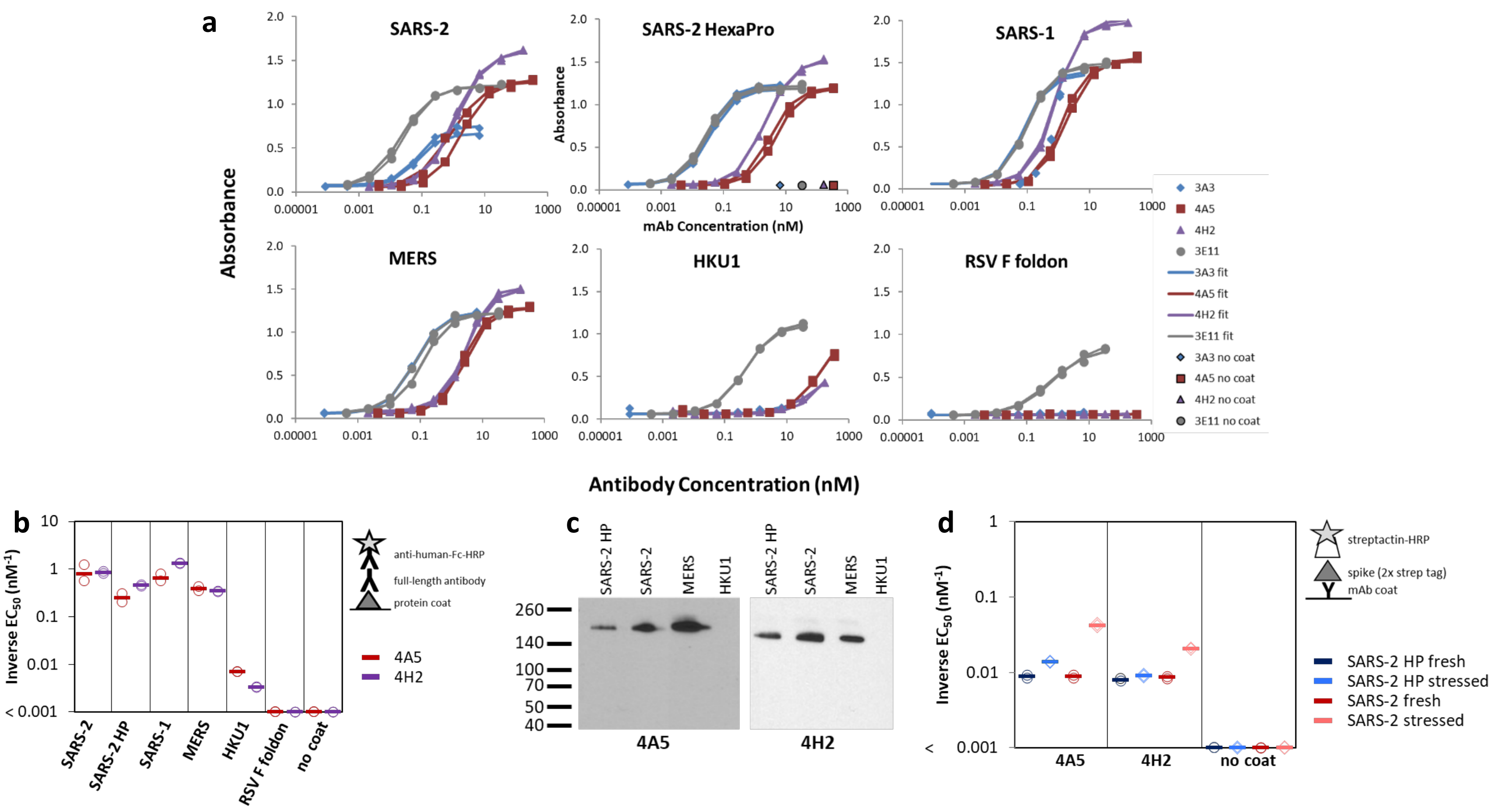


**Supplementary Figure 5. Antibodies 3A3, 4A5, and 4H2 bind highly pathogenic coronaviruses. a** SARS-CoV-2 spike (SARS-2), SARS-CoV-2 HexaPro spike (SARS-2 HP), SARS-CoV spike (SARS-1), MERS-CoV spike, and HKU1 spike, RSV F, and milk (no coat) were coated on high binding plates for ELISA. Full length antibodies were serially diluted in duplicate and allowed to bind the coat proteins, then binding was detected with anti-human Fc-HRP development of TMB substrate. The binding data and antibody concentrations for each replicate were fit to a four-parameter logistic curve. Inverse $EC_{50}$ of these data are presented in Fig. 1a and in panel **b** of this Supplementary Figure. This experiment was repeated twice; data shown are representative of replicate experiments. **c** Spike variants were western blotted with 4A5 and 4H2. **d** Heat and freeze thaw stressed SARS-2 or SARS-2 HP spike was captured by 4A5 and 4H2 antibodies.

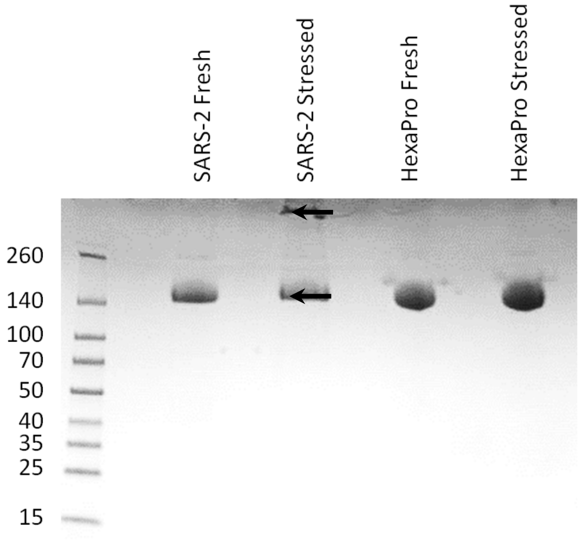


**Supplementary Figure 6. SDS-PAGE analysis of fresh and stressed SARS-CoV-2 and SARS-CoV-2 HexaPro spikes shows a substantial aggregation of the stressed SARS-CoV-2 spike.** For each spike, 8 μg of protein was analyzed by SDS-PAGE under non-reducing conditions. Reduced band intensity of stressed SARS-CoV-2 spike (bottom arrow) and appearance of a band just under the loading wells (top arrow) indicates aggregation products not apparent in the other samples. Densitometry analysis (ImageJ) of these bands indicates that the stressed SARS-CoV-2 spike's main peak is diminished by ~20% relative to the fresh SARS-CoV-2 spike intensity. Ladder molecular weights in kDa are indicated to the left of the gel image.

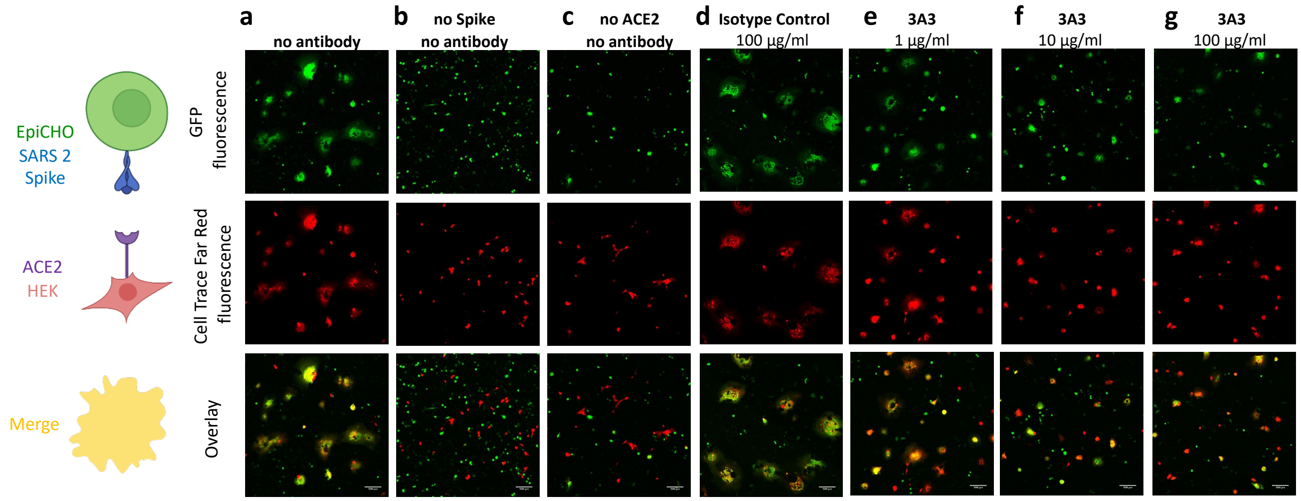


**Supplementary Figure 7. 3A3 inhibits cellular fusion induced by the interaction of SARS-CoV-2 spike with human ACE2. a** HEK 293 cells stably expressing human ACE2 were stained with Cell Trace Far Red and incubated with a CHO-based cell line transiently expressing wild-type SARS-CoV-2 spike and EGFP. The cultures were imaged after 24 hours of incubation for EGFP (green) or Cell Trace Far Red (red) and the level of colocalization (yellow) was evaluated. **b** CHO cells not expressing SARS-CoV-2 spike and **c** HEK 293 cells not expressing ACE2 exhibited minimal fusion. **d** When the cultures were preincubated with an irrelevant isotype control antibody, extensive fusion and syncytia formation equivalent to no antibody was apparent. Incubation at **e** 6.7, **f** 67, and **g** 670 nM 3A3 reduced fusion in a dose-dependent manner with significance reached at 67 nM. Scale bar, 100 µm.

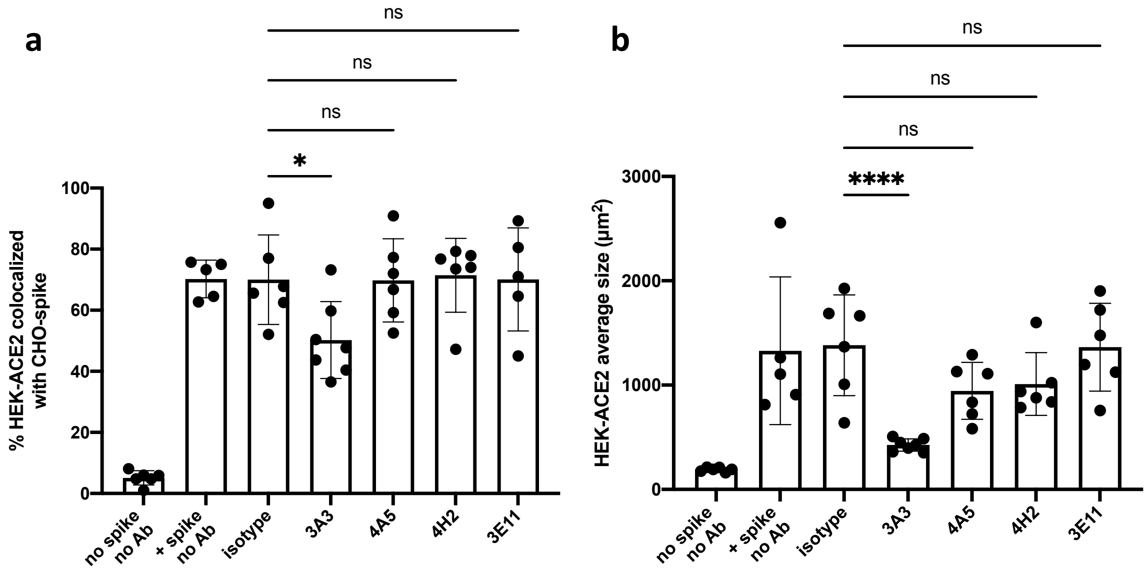


**Supplementary Figure 8. Antibodies 4A5, 4H2, and 3E11 have no impact on cellular fusion.** HEK 293 cells stably expressing ACE2 were stained with Cell Trace Far Red and mixed with CHO cells expressing only EGFP (GFP only) or CHO cells expressing EGFP and wild-type SARS-CoV-2 spike and preincubated with 200 nM of: no antibody, isotype control, 3A3, 4H2, 4A5, or 3E11. After evaluating **a**, the percentage of cells with colocalized red and green fluorescence and **b** the average HEK-ACE2 cell size, only incubation with 3A3 was found to reduce fusion significantly. Shown are the mean and standard deviation of at least 100 cells per condition from 5-7 independent images. The statistical analysis was performed with ANOVA. Results show are representative of three independent experiments; * $p < 0.05$, **** $p < 0.0001$.

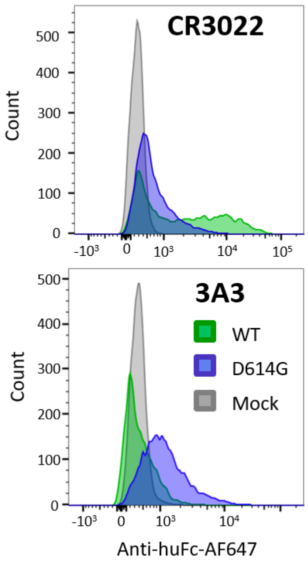


**Supplementary Figure 9. The 3A3 epitope is accessible on unmodified SARS-CoV-2 spike.** Antibody 3A3 weakly stains wild-type SARS-CoV-2 spike (WT; green) expressing Expi293 cells and more strongly binds SARS-CoV-2 D614G spike (blue) expressing cells in flow cytometry. Control Expi293 cells not expressing spike (mock) are shown in grey.

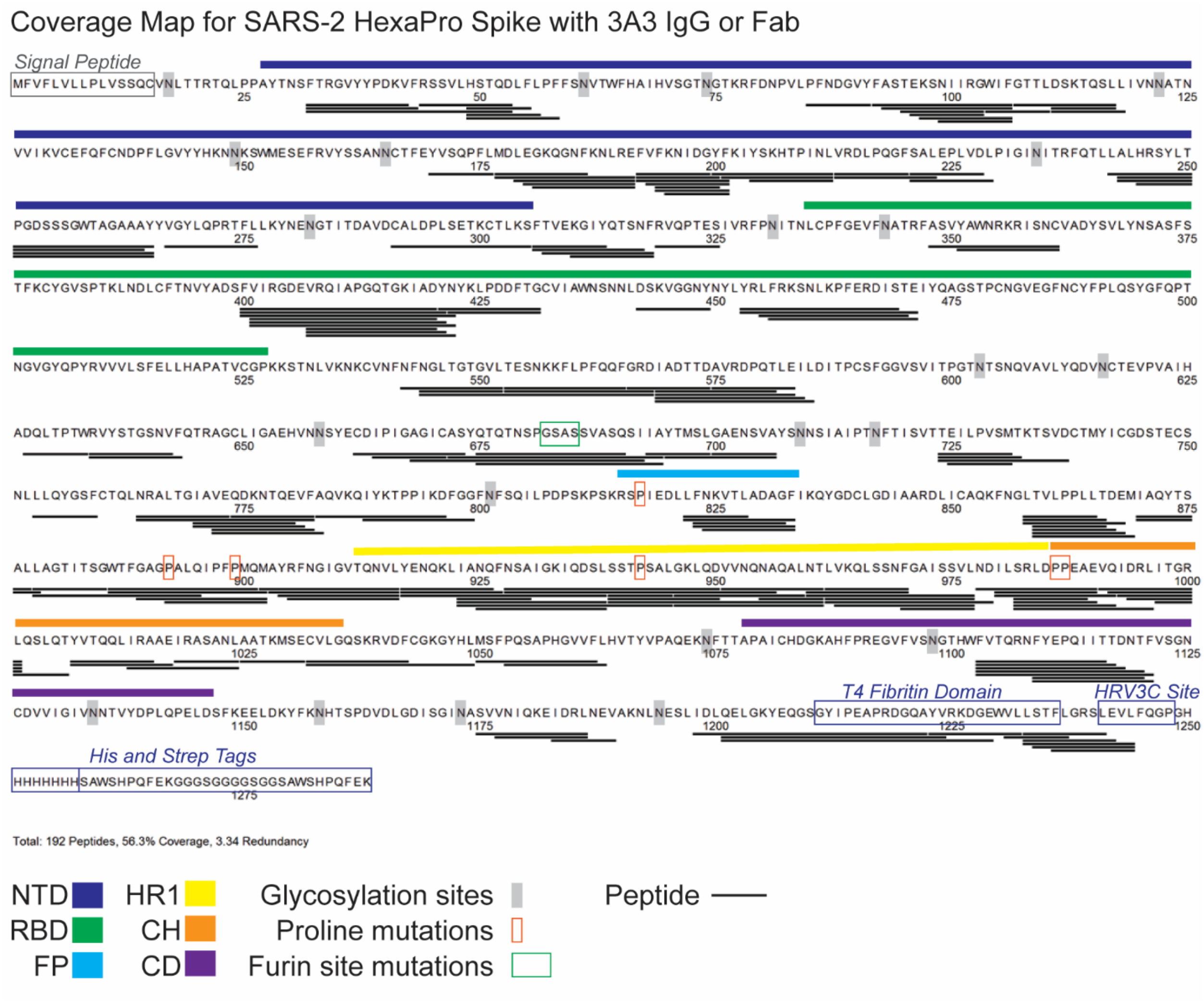


**Supplementary Figure 10. Peptides monitored through all timepoints of deuteration for SARS-CoV-2 HexaPro spike alone and with 3A3 IgG or Fab.** A total of 192 peptides were monitored, covering 56.3% of the SARS-CoV-2 HexaPro spike sequence and averaging 3.34 redundancy per amino acid. All peptides were manually checked. SARS-CoV-2 spike features are indicated on the sequence, including glycosylation sites. Glycosylation was not included in the peptide search and so no peptides are recovered surrounding these sites.

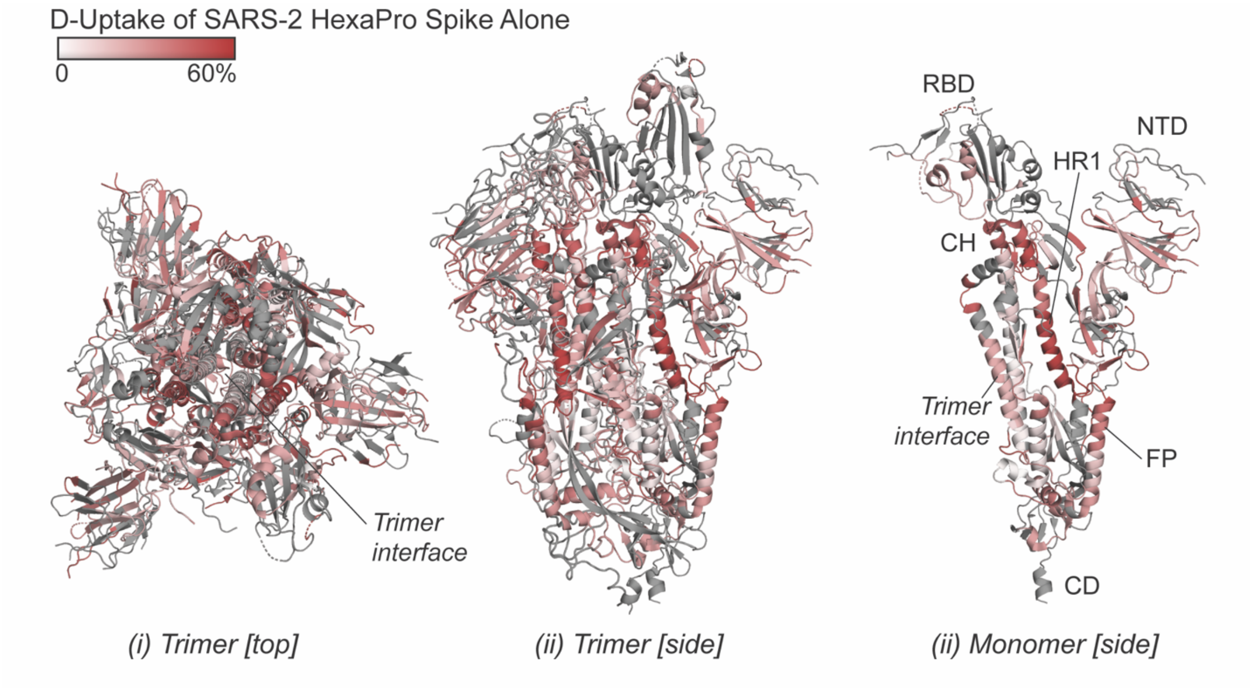


**Supplementary Figure 11. Deuterium uptake of SARS-CoV-2 HexaPro spike alone suggests the trimer is maintained under HDX conditions.** Trimeric (i and ii) and monomeric (iii) SARS-CoV-2 2P spike (PDB: 6VSB) colored according to fractional deuterium uptake of the SARS-CoV-2 HexaPro spike alone after $10^3$ s of exchange. The figure was prepared using DynamX per residue output and Pymol. Residues lacking coverage are indicated in grey. Structural features are labeled, including the central trimer interface that shows relatively low deuterium incorporation.

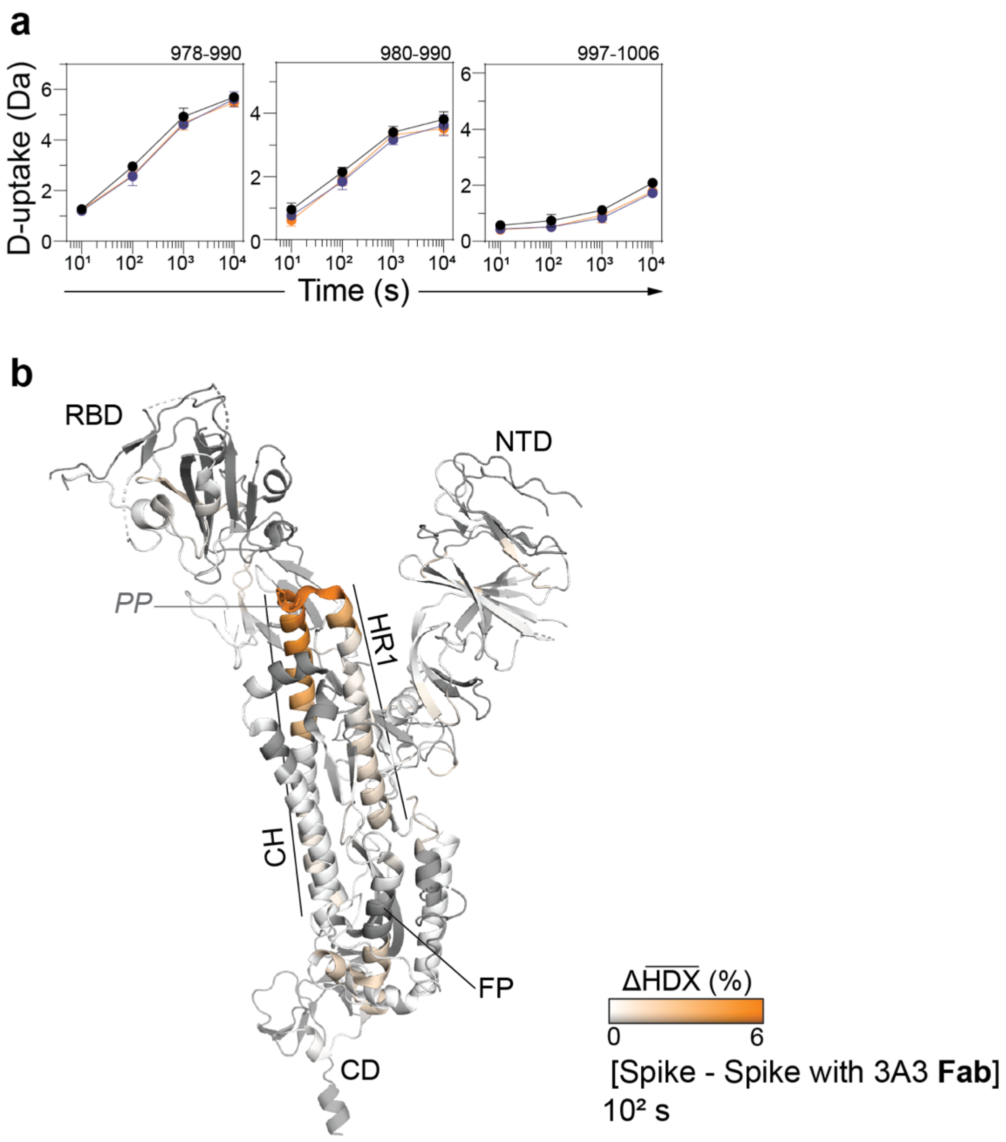


**Supplementary Figure 12. Location of the 3A3 epitope in SARS-CoV-2 spike a** Additional deuterium uptake plots for peptides with a significant decrease in deuterium uptake upon addition of 3A3 (see also Fig. 4C). Traces are SARS-CoV-2 HexaPro spike alone (black), with 3A3 IgG (blue), and with 3A3 Fab (orange). Error bars are ±2σ from 3 or 4 technical replicates. Y-axis is 70% of max deuterium uptake assuming the N-terminal residue undergoes complete back-exchange. Data have not been corrected for back-exchange. **b** Monomeric SARS-CoV-2 2P spike (PDB: 6VSB chain B) colored according to the difference in deuterium fractional uptake between SARS-CoV-2 HexaPro spike alone and with 3A3 Fab at $10^2$ s. The figure was prepared using DynamX per residue output without statistics and Pymol. Residues lacking coverage are indicated in grey. Structural features are labeled, including the 2P mutations at residues 986 and 987 (shown as sticks).

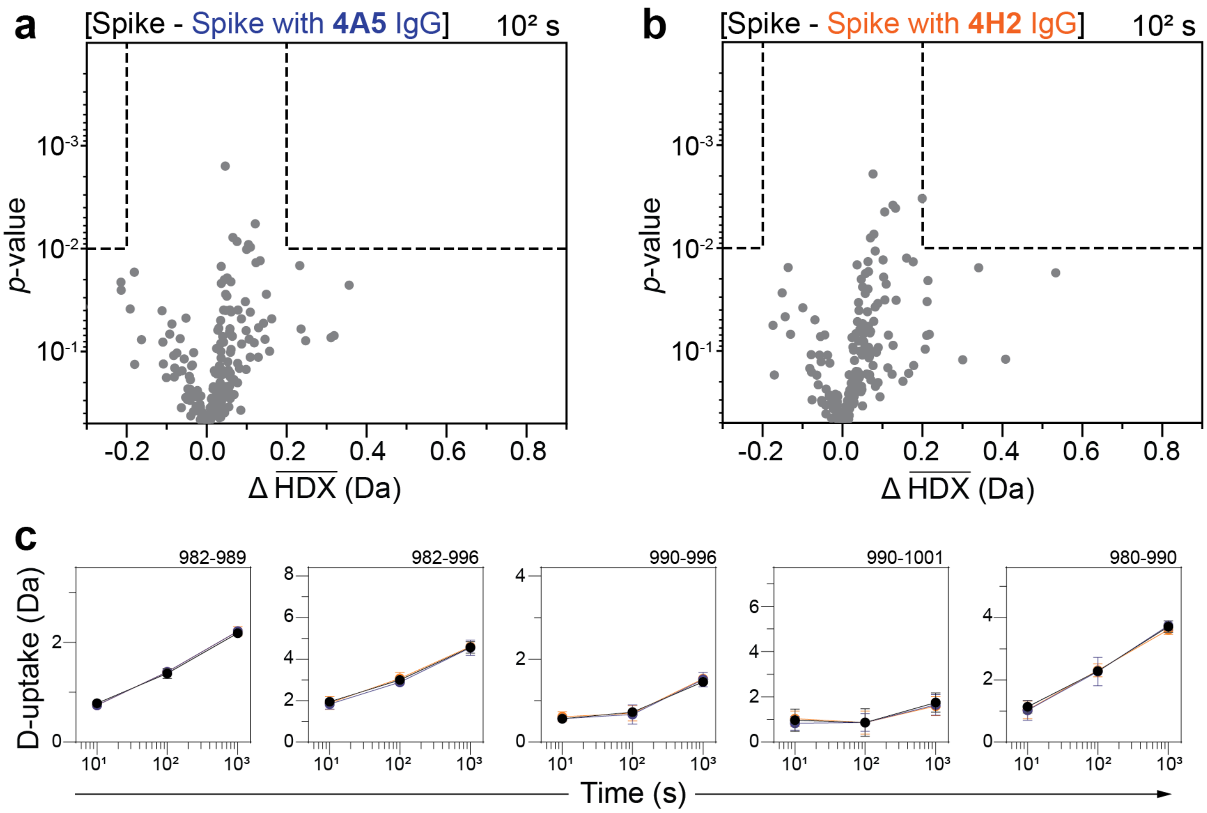


**Supplementary Figure 13. 4A5 and 4H2 do not bind the same epitope as 3A3.** Volcano plots showing changes in deuterium uptake in SARS-CoV-2 HexaPro spike peptides upon addition of 4A5 **a** or 4H2 **b** after $10^2$ s exchange. Significance cutoffs are an average change in deuterium uptake greater than 0.2 Da and a *p*-value less than $10^{-2}$ in a Welch's t-test (hatched box). **c** Deuterium uptake plots for peptides from the 3A3 epitope. Traces are SARS-CoV-2 HexaPro spike alone (black), with 4A5 (blue), and with 4H2 (orange). Error bars are ±2σ from 4 technical replicates. Y-axis is 70% of max deuterium uptake assuming the N-terminal residue undergoes complete back-exchange. Data have not been corrected for back-exchange—all traces overlay.

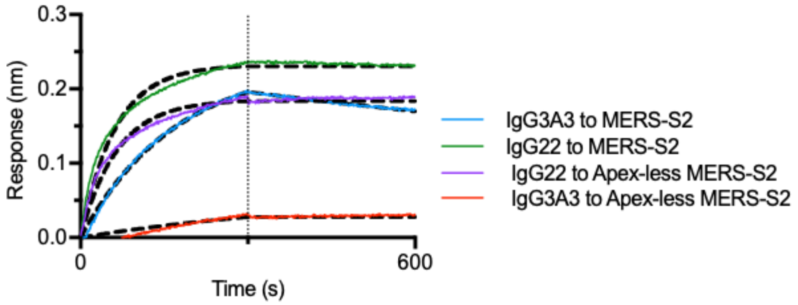


**Supplementary Figure 14. Antibody 3A3 cannot bind the MERS S2 domain with the apex removed.** To evaluate 3A3 binding to MERS S2 with the apex region containing the 3A3 epitope removed by BLI, anti-human Fc Sensors were used to pick up 3A3 (10 nM) or an S2 binding control antibody IgG22 to a response of 0.6 nm. Then mAb coated tips were dipped into wells containing MERS-S2 or apex-less MERS-S2 (100 nM). Data were collected over an antibody association to the MERS S2 domains for 5 minutes and dissociation for 5 minutes. The colored line shows data collected; the black dashed line shows the best fit.

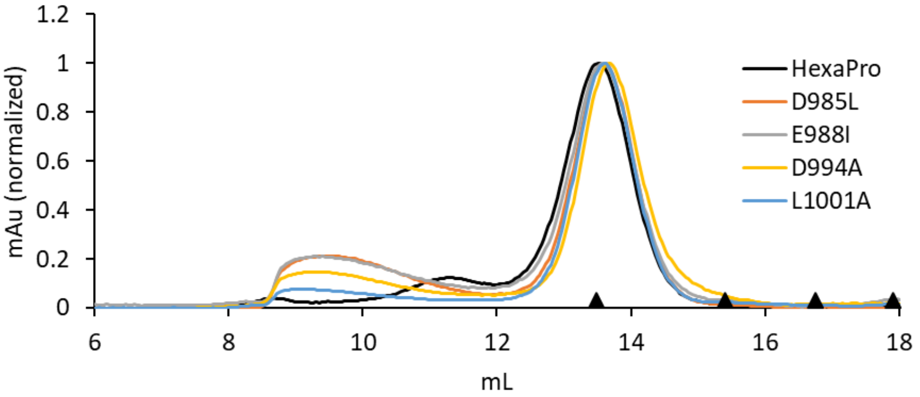


**Supplementary Figure 15. HexaPro variants with reduced 3A3 binding retain trimer SEC profile.** The single point mutants of HexaPro, D985L, E988I, D994A, and L1001A, had reduced binding to 3A3 relative to HexaPro (Fig. 4c), but retained the overall size of unmodified HexaPro by SEC with elution of the main peak at ~13.75 mL on a Superose 6 Increase 30/100 column, indicating the spike was intact. Molecular weight markers (black triangles) on the x-axis are peak elution volumes from the following standards in order from left to right: thyroglobulin (669 kDa, 13.49 mL), ferritin (440 kDa, 15.40 mL), β-amylase (200 kDa, 16.75 mL), aldolase (158 kDa, 17.91 mL)

**a**

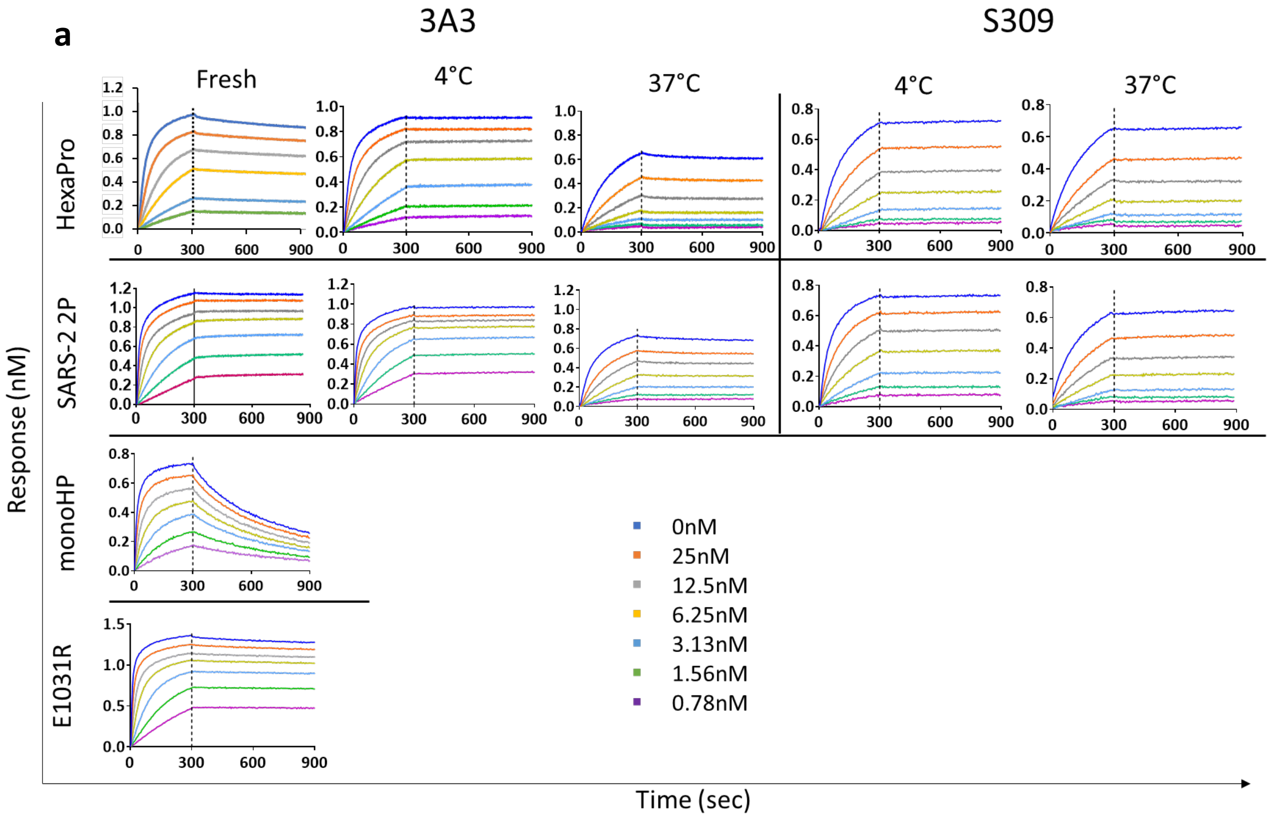


**b**

| | 3A3 | | | | | | | | S309 | | | |
|---|---|---|---|---|---|---|---|---|---|---|---|---|
| | monoHP | E1031R | HexaPro | | | 2P | | | HexaPro | | 2P | |
| | Fresh | Fresh | Fresh | 4°C | 37°C | Fresh | 4°C | 37°C | 4°C | 37°C | 4°C | 37°C |
| $K_d$ (nM) | 3.6 ± 0.7 | 1.5 ± 0.1 | 10 ± 3 | 8 ± 3 | 30 ± 20 | 2.7 ± 0.3 | 1.9 ± 0.2 | 12 ± 1 | 18 ± 2 | 24 ± 2 | 7 ± 5 | 20 ± 1 |
| $k_{on}$ ($\mu M^{-1}s^{-1}$) | 2.70 ± 0.04 | 2.88 ± 0.07 | 0.74 ± 0.04 | 0.58 ± 0.02 | 0.16 ± 0.02 | 1.51 ± 0.02 | 1.4 ± 0.2 | 0.30 ± 0.02 | 0.21 ± 0.02 | 0.18 ± 0.02 | 0.39 ± 0.02 | 0.21 ± 0.03 |

**Supplementary Figure 16. Binding of 3A3 to spike is impacted by accessibility to the epitope. a** Binding of 3A3 and S309 to SARS-CoV-2 spikes were evaluated by BLI. Full length antibodies were captured by anti-human Fc sensors, dipped into serial dilutions of spike from 25 to 0.78 nM and then incubated in buffer. SARS-CoV-2 S2P or HexaPro spike was freshly thawed from preparations immediately stored at -80°C (fresh), stored at 4°C for 1-3 weeks (4°C), or stored at 37°C for 24 hours (37°C) before dilution into buffer immediately before BLI analysis. Vertical lines indicate the start of the dissociation phase. Off-rates with a trimeric analyte (HexaPro, SARS-CoV-2, and E1031R spikes) are not reliable due to rebinding. **b** Equilibrium $K_d$ and on-rates were calculated based on seven concentrations in two independent experiments (representative data shown in **a**). For all fits the coefficient of determination ($R^2$) was >0.98 and $R_{max}$ was between 0.7 and 1.4 nM.

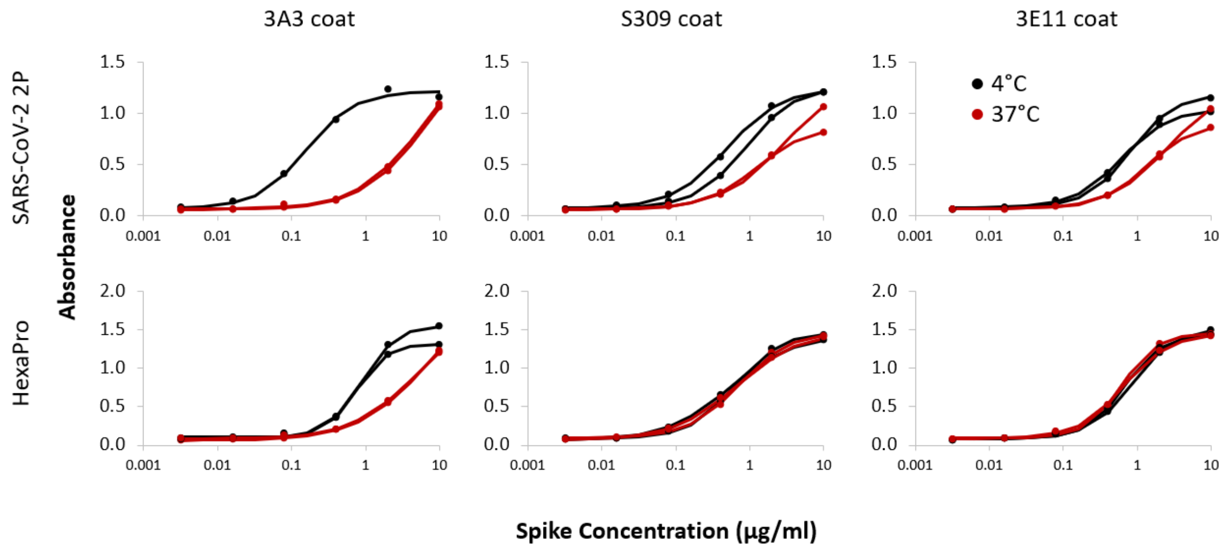


**Supplementary Figure 17. Epitope accessibility affects antibody binding by ELISA.** SARS-CoV-2 S2P or HexaPro spike stored at 4°C for 1-3 weeks (4°C, black), or stored at 37°C for 24 hours (37°C, red) were allowed to bind to 3A3, S309, or 3E11 coated plates for one hour at room temperature. Detection with StrepTactin-HRP and TMB substrate showed reduced binding of 3A3 after incubation at 37°C. Control antibodies S309 and 3E11 had somewhat reduced binding to SARS-CoV-2 spike incubated at 37°C likely due to protein degradation, aggregation, or tag loss.

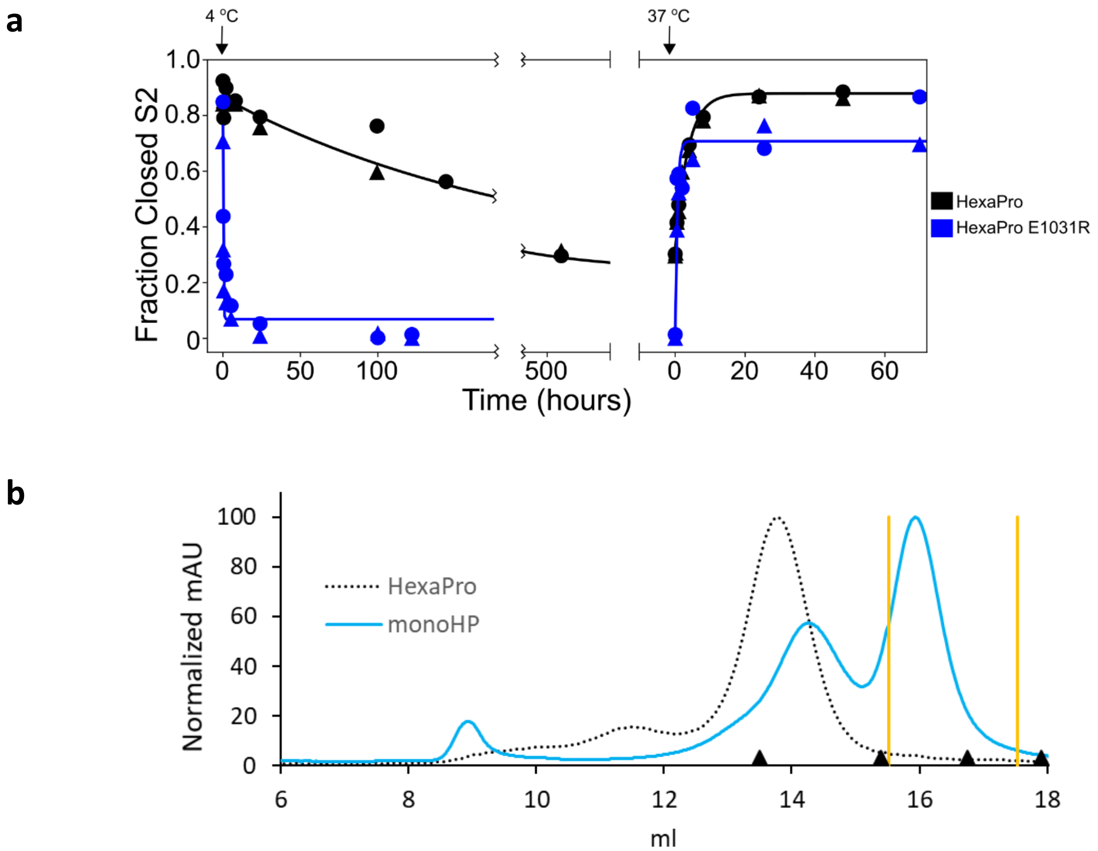


**Supplementary Figure 18. HexaPro variants expose the 3A3 epitope. a** The kinetics of interconversion between the closed-S2 and open-S2 states of HexaPro and HexaPro E1031R were evaluated by HDX-MS as previously described (Costello, Shoemaker, et al 2021). The spike proteins were incubated at 37°C for 24 hours, then incubated at 4°C, assayed for detection of the S2-open state over time, then transferred to 37°C and assayed again. The fraction closed-S2 was estimated by exposing a sample of the incubation reaction to a 1 min pulse of deuterium and quenching the labeling reaction with low pH and temperature. After LC-MS and peptide identification, the bimodal mass envelopes for all timepoints for one peptide were globally fit to a sum of two gaussians, keeping the center and width of each gaussian constant across all incubation time points. After fitting, the area under the lower molecular weight gaussian was integrated to determine the fraction of closed-S2. For HexaPro, the half-life of conversion from closed to open (37°C → 4°C was 143.5 h and the half-life of conversion from open to closed (4°C → 37°C) was 2.5 h. These conversions for HexaPro E1031R occur with half-lives decreased by >700-fold (0.2 h) and >6-fold (0.4 h), respectively. **b** An HRV3C cut site was inserted between the HexaPro spike c-terminus and the foldon domain allowing expression of the spike as a trimer, incubation at 4°C for 5 days, then digestion with HRV3C protease to produce HexaPro monomer (monoHP). After digestion, the sample was purified by SEC on a Superose6 column to isolate monoHP (blue line) from partially digested products. Fractions between the yellow lines were collected for use in BLI. Molecular weight markers (black triangles) on the x-axis are peak elution volumes from the following standards in order from left to right: thyroglobulin (669 kDa, 13.49 mL), ferritin (440 kDa, 15.40 mL), β-amylase (200 kDa, 16.75 mL), aldolase (158 kDa, 17.91 mL)

**Supplementary Table 1. HDX summary table**

| Data Set | SARS-2 HexaPro spike | SARS-2 HexaPro spike + 3A3 IgG | SARS-2 HexaPro spike + 3A3 Fab | SARS-2 HexaPro spike | SARS-2 HexaPro spike + 4A5 IgG | SARS-2 HexaPro spike + 4H2 IgG |
|---|---|---|---|---|---|---|
| **HDX reaction details** | 200 mM NaCl, 20 mM Tris | | | 200 mM NaCl, 20 mM Tris | | |
| | 0.5 uM S2 | 0.5 uM S2 + 0.55 uM 3A3 IgG | 0.5 uM S2 + 0.55 uM 3A3 Fab | 0.5 uM S2 | 0.5 uM S2 + 0.75 uM 4A5 IgG | 0.5 uM S2 + 0.75 uM 4H2 IgG |
| | pHread = 7.6 | | | pHread = 7.6 | | |
| **HDX time course (s)** | 10, 100, 1000, 10000 at 25°C | | | 10, 100, 1000, 10000 at 25°C | | |
| **HDX control samples** | Unlabeled S2 | | | Unlabeled S2 | | |
| **Back-exchange (mean)** | ~40% | | | ~40% | | |
| **# of peptides** | 192 | | | 188 | | |
| **Sequence coverage** | 56.30% | | | 60.00% | | |
| **Average Peptide length / Redundancy** | 13/3.34 | | | 13/3.21 | | |
| **Replicates (biological or technical)** | 4 (technical) | | | 4 (technical) | | |
| **Repeatability** | 0.10 Da (average standard deviation) | | | 0.087 Da (average standard deviation) | | |
| **Significant difference** | Average ΔHDX greater than 0.2 Da and p-value less than 0.01 | | | Average ΔHDX greater than 0.2 Da and p-value less than 0.01 | | |

**Supplementary Table 2. Complete HDX data attached as an excel file.**

**Supplementary Table 3. Width of Isotopic Distributions for Example Spike Peptides.** Peptides were selected from regions identified as being bimodal in *Costello et al*. We had no coverage in residues 626-636 and 1146-1166. Peak width (PW) in Da was calculated in triplicate for each peptide in the non-deuterated sample (ND Control) and at each time point of exchange. The SD is reported. The change in peak width (ΔPW) was calculated by subtracting the PW of the control from the PW of the sample. Bimodality was assessed by taking the maximum peak width for a particular peptide ($\Delta PW_{max}$) and assessing if it was greater than 2 Da. Peak width was calculated using a method similar to Weis et al. Peptides were centroided with the Apex3D algorithm using DynamX (Waters). Following manual curation, ion stick data were transferred into Excel as two columns of data, m/z values and intensities, and the maximum peak in the isotopic envelope was determined. The list was then searched in descending m/z order to identify the two lowest m/z peaks that straddled 20% of the maximum peak's intensity. The m/z value at an envelope intensity of 20% of the maximum intensity was determined using linear interpolation between these two peaks. This process was repeated with a search in ascending m/z order. The relative peak width was determined by multiplying by z (the charge state). For peptide spectra without peaks straddling 20% of the maximum peak's intensity on one sides of the maximum, typical for lower m/z peaks for peptides exhibiting low deuteration, the farthest isotopic centroid peak on that side was used as the m/z limit for calculating peak width while the other m/z limit was determined using the previously described method.

| **Peptide** | | | | **ND Control** | | **10 s HDX** | | | **$10^2$ s HDX** | | | **$10^3$ s HDX** | | | **$10^4$ s HDX** | | | **Possible Bimodality** | |
|---|---|---|---|---|---|---|---|---|---|---|---|---|---|---|---|---|---|---|---|
| **Start** | **End** | **Sequence** | **Length** | **PW (Da)** | ***SD*** | **PW (Da)** | ***SD*** | **ΔPW (Da)** | **PW (Da)** | ***SD*** | **ΔPW (Da)** | **PW (Da)** | ***SD*** | **ΔPW (Da)** | **PW (Da)** | ***SD*** | **ΔPW (Da)** | **$\Delta PW_{max}$ (Da)** | **$\Delta PW_{max}$ > 2 Da** |
| 291 | 305 | CALDPLSETKCTLKS | 15 | 2.8 | *0.1* | 4.7 | *0.2* | **1.9** | 4.7 | *0.0* | **1.9** | 5.4 | *0.1* | **2.6** | 5.5 | *0.3* | **2.7** | **2.7** | ✓ |
| 553 | 565 | TESNKKFLPFQQF | 13 | 2.8 | *0.0* | 7.0 | *0.1* | **4.1** | 7.0 | *0.2* | **4.1** | 6.2 | *0.3* | **3.3** | 6.2 | *0.4* | **3.4** | **4.1** | ✓ |
| 553 | 568 | TESNKKFLPFQQFGRD | 16 | 3.5 | *0.1* | 8.5 | *0.1* | **5.1** | 8.7 | *0.1* | **5.2** | 7.3 | *0.2* | **3.8** | 7.2 | *0.1* | **3.7** | **5.2** | ✓ |
| 634 | 642 | RVYSTGSNV | 9 | 1.8 | *0.1* | 5.9 | *0.3* | **4.1** | 5.9 | *0.1* | **4.1** | 5.5 | *0.2* | **3.7** | 5.8 | *0.2* | **4.0** | **4.1** | ✓ |
| 634 | 643 | RVYSTGSNVF | 10 | 1.9 | *0.1* | 6.1 | *0.0* | **4.2** | 5.9 | *0.1* | **4.0** | 6.1 | *0.1* | **4.2** | 5.7 | *0.4* | **3.8** | **4.2** | ✓ |
| 668 | 692 | AGICASYQTQTNSPGSASSVASQSI | 25 | 3.4 | *0.1* | 7.4 | *0.3* | **3.9** | 8.1 | *0.4* | **4.7** | 7.9 | *0.1* | **4.5** | 8.0 | *0.2* | **4.6** | **4.7** | ✓ |
| 870 | 878 | IAQYTSALL | 9 | 1.8 | *0.0* | 2.6 | *0.2* | **0.9** | 2.7 | *0.2* | **1.0** | 2.9 | *0.1* | **1.1** | 2.8 | *0.1* | **1.0** | **1.1** | × |
| 902 | 916 | MAYRFNGIGVTQNVL | 15 | 3.1 | *0.2* | 5.8 | *0.1* | **2.6** | 6.4 | *0.3* | **3.3** | 6.9 | *0.1* | **3.8** | 7.6 | *0.3* | **4.4** | **4.4** | ✓ |
| 980 | 990 | ILSRLDPPEAEVQ | 11 | 2.1 | *0.1* | 4.0 | *0.2* | **1.8** | 4.7 | *0.1* | **2.6** | 4.9 | *0.1* | **2.7** | 5.4 | *0.3* | **3.2** | **3.2** | ✓ |
| 980 | 992 | ILSRLDPPEAEVQ | 13 | 2.4 | *0.1* | 4.5 | *0.3* | **2.2** | 6.2 | *0.2* | **3.8** | 6.2 | *0.1* | **3.8** | 6.0 | *0.2* | **3.7** | **3.8** | ✓ |
| 982 | 989 | SRLDPPEA | 8 | 1.7 | *0.1* | 3.0 | *0.1* | **1.3** | 3.9 | *0.1* | **2.2** | 3.7 | *0.1* | **2.0** | 3.9 | *0.0* | **2.2** | **2.2** | ✓ |
| 982 | 992 | SRLDPPEAEVQ | 11 | 2.6 | *0.1* | 4.9 | *0.1* | **2.3** | 5.2 | *0.4* | **2.6** | 5.4 | *0.1* | **2.9** | 5.4 | *0.1* | **2.9** | **2.9** | ✓ |
| 991 | 1001 | VQIDRLITGRL | 11 | 2.7 | *0.1* | 4.3 | *0.4* | **1.5** | 4.2 | *0.2* | **1.4** | 5.1 | *0.4* | **2.3** | 7.3 | *0.1* | **4.5** | **4.5** | ✓ |
| 992 | 1001 | QIDRLITGRL | 10 | 2.2 | *0.2* | 3.5 | *0.4* | **1.3** | 3.3 | *0.3* | **1.1** | 4.0 | *0.2* | **1.8** | 5.9 | *0.4* | **3.7** | **3.7** | ✓ |
| 997 | 1006 | ITGRLQSLQT | 10 | 1.9 | *0.0* | 3.0 | *0.0* | **1.2** | 3.4 | *0.2* | **1.5** | 3.8 | *0.1* | **1.9** | 4.8 | *0.1* | **3.0** | **3.0** | ✓ |
| 1007 | 1015 | YVTQQLIRA | 9 | 2.8 | *0.1* | 3.7 | *0.1* | **0.9** | 3.8 | *0.1* | **1.0** | 4.0 | *0.1* | **1.2** | 4.6 | *0.2* | **1.8** | **1.8** | × |
| 1013 | 1024 | IRAAEIRASANL | 12 | 2.3 | *0.1* | 3.5 | *0.2* | **1.2** | 3.5 | *0.3* | **1.2** | 4.3 | *0.1* | **2.0** | 5.4 | *0.4* | **3.1** | **3.1** | ✓ |
| 1018 | 1024 | IRASANL | 7 | 1.8 | *0.1* | 2.5 | *0.1* | **0.8** | 2.5 | *0.2* | **0.8** | 2.8 | *0.1* | **1.0** | 3.6 | *0.1* | **1.8** | **1.8** | × |
| 1183 | 1189 | IDRLNEV | 7 | 1.6 | *0.1* | 3.1 | *0.1* | **1.5** | 3.6 | *0.2* | **1.9** | 3.7 | *0.1* | **2.1** | 3.9 | *0.1* | **2.3** | **2.3** | ✓ |
